# Divalent cations govern the activity of coronavirus nsp15

**DOI:** 10.1101/2025.03.29.646087

**Authors:** Xionglue Wang, Jing Li, Zhichao Liu, Longfei Wang, Bin Zhu

**Affiliations:** Key Laboratory of Molecular Biophysics, the Ministry of Education, College of Life Science and Technology, Huazhong University of Science and Technology, Wuhan, Hubei 430074, China; Department of Cardiology, Taikang Center for Life and Medical Sciences, Zhongnan Hospital of Wuhan University, School of Pharmaceutical Sciences, Wuhan University, Wuhan, Hubei 430074, China; Shenzhen Huazhong University of Science and Technology Research Institute, Shenzhen 518063, China

**Author notes:** These authors contributed equally (Xionglue Wang, Jing Li). Correspondence (X.L.W.), (L.F.W.), (B.Z.).

## Abstract

Coronavirus nsp15 is an endoribonuclease that limits the accumulation of viral dsRNA in cytosol and plays a crucial role in the evasion of host immunity. Here, we show that Co^2+^ or Ni^2+^ potently activates nsp15 of human and animal coronaviruses, including SARS-CoV-2, SARS-CoV, MERS-CoV, HCoV-229E, MHV-A59, and PEDV. Whereas Zn^2+^ strongly inhibits nsp15 of these coronaviruses. Cryo-EM structures of the Co^2+^-bound SARS-CoV-2 nsp15/dsRNA complexes reveal that Co^2+^ binds to the nsp15 active site and is coordinated by three catalytic residues H234, H249, K289, and the 2’ hydroxyl of the flipped base from dsRNA, suggesting that Co^2+^ stabilizes the catalytic core and increases substrate binding. Although our data indicated that Ni^2+^ and Zn^2+^ bind to the same site as Co^2+^, Zn^2+^ lacks the catalytic capability and inhibits nsp15. These findings reveal that divalent cations govern coronavirus nsp15’s activity and provide basis for new therapeutic strategies.

## INTRODUCTION

Coronaviruses (CoVs) are a diverse class of single-stranded (ss) positive-sense RNA viruses that infect humans, other mammals and avian species. Since the beginning of the 21st century, three distinct coronaviruses (severe acute respiratory syndrome (SARS)-CoV, Middle East respiratory syndrome (MERS)-CoV, and SARS-CoV-2) have emerged that are capable of causing severe disease in humans.^1^ Hosts generally combat viruses via interferon (IFN) responses and chemokine-mediated leukocyte recruitment.^2,3^ However, SARS-CoV-2 and SARS-CoV evade IFN responses, enabling persistent replication while triggering excessive chemokine production, causing dysregulated immunity and disease.^4,5^ It is noteworthy that the fifteenth nonstructural protein (nsp15) encoded by the coronavirus genome plays a crucial role in counteracting host IFN responses.^6^

The replication and transcription of the coronaviral positive-stranded RNA genome inevitably produces double-stranded RNA (dsRNA) as an intermediate.^1^ DsRNA is known as an important pathogen-associated molecular pattern, capable of triggering IFN responses through the activation of dsRNA sensors.^7^ The C-terminal domain of coronavirus nsp15 belongs to the uracil-specific endoribonuclease (EndoU) family,^8^ which enables nsp15 to cleave the 3’-side of uridylate in RNA.^9^ It is widely accepted that the EndoU activity of coronavirus nsp15 including SARS-CoV-2 nsp15 plays a role in evading host dsRNA sensors by limiting the accumulation of viral dsRNA in the cytoplasm.^10–13^ Nevertheless, it has been proposed that nsp15 could antagonize IFN responses through other strategies.^14–16^ In our previous study, we demonstrated that SARS-CoV-2 nsp15 preferentially cleaves AU-rich dsRNA and found that coronavirus genomes generally have a high AU content, with an average value of 61%.^17^ Therefore, we hypothesized that after processing by nsp15, the degradation products of viral dsRNA intermediates are not able to efficiently activate dsRNA sensors. In addition, two studies on mouse hepatitis virus (MHV) nsp15 proposed two potential mechanisms for reducing viral dsRNA, based on cleavage of viral positive or negative-stranded ssRNA.^18,19^ It is important to note that, given its co-localization with the viral replication and transcription complex (RTC) and replicating viral RNA,^10,20–22^ the EndoU activity of nsp15 must be regulated in an ordered manner to ensure that viral RNA is not degraded by nsp15 during genome replication and transcription, and that residual viral dsRNA is degraded by nsp15 efficiently following the completion of genome replication and transcription or when the viral RNA synthesis is stalled. However, the regulatory mechanism of nsp15’s activity is poorly understood.

Mn^2+^ was found to be able to enhance the *in vitro* RNA cleavage activity of nsp15.^23–26^ However, the enhancement was reported to be conditional and substrate-dependent,^27,28^ and usually observed at millimolar concentrations of Mn^2+^, which is physiologically irrelevant.^29,30^ In addition, the effect of Mn^2+^ on nsp15 stability and oligomerization was proved to be subtle,^17,27,31–33^ and all attempts to identify the presence of Mn^2+^ in the nsp15 structure have been unsuccessful.^26,34,35^ These findings suggest that the interaction between Mn^2+^ and nsp15 may be non-specific or non-structural.

Here, we discovered that both Co²⁺ and Ni²⁺ exhibit a strong ability to activate nsp15, with this activation being significantly more potent than that triggered by Mn²⁺. On the other hand, Zn²⁺ was found to have a pronounced inhibitory effect on nsp15’s activity. We determined cryo-electron microscopy (Cryo-EM) structures of the Co^2+^-bound SARS-CoV-2 nsp15/dsRNA complexes, which reveal that Co^2+^ binds to the nsp15 active site and is coordinated by three catalytic residues H234, H249, K289, and the 2’ hydroxyl of the flipped base from dsRNA, suggesting that Co^2+^ stabilizes the catalytic core and increases substrate binding. The results for the H249A mutant suggest that Ni^2+^ and Zn^2+^ also act through the active site. These findings reveal that divalent cations govern coronavirus nsp15’s activity and offered insights into antiviral therapeutic strategies.

## RESULTS

### Co^2+^ and Ni^2+^ strongly activate SARS-CoV-2 nsp15

We initially investigated the impact of divalent metal ions on SARS-CoV-2 nsp15 in buffers containing dithiothreitol (DTT), which had usually been used for nsp15 cleavage reactions in previous studies. However, both Co^2+^ and Ni^2+^ appeared to undergo reduction in buffers containing DTT, as evidenced by a color change from clear to red (Figure S1A). This limited the examination of the role of Co^2+^ and Ni^2+^ on nsp15 (Figure S1B). Therefore, we decided to employ a buffer without DTT. To this end, we conducted a comprehensive investigation into the impact of DTT on nsp15 activity. The results showed that DTT had no significant effect on the RNA cleavage efficiency, RNA cleavage preference and protein stability of nsp15 (Figure S1C–S1G). The cleavage reactions were conducted with Co^2+^ and Ni^2+^ under conditions lacking DTT, with the unexpected results that Co^2+^ and Ni^2+^ significantly activated nsp15, to a much greater extent than Mn^2+^ (Figure S1B). To gain further insight into this activation, an ssRNA substrate of 44 nt (ssRNA44) and a dsRNA substrate of 44 bp (dsRNA44), which are identical in sequence, were employed (Figure 1A). Our previous research had already demonstrated that under Mn^2+^ conditions, nsp15 exhibited a preferential cleavage of consecutive Us located in an AU-rich area in dsRNA44, and that nsp15 cleaved almost all U sites in ssRNA44, although the cleavage efficiencies at these sites exhibited slight variations. We performed the titration experiments with varying concentrations of Mn^2+^, Co^2+^ and Ni^2+^. The results indicated that Co^2+^ and Ni^2+^ activated nsp15 much more strongly than Mn^2+^, without a notable alteration in nsp15 cleavage preference (Figure 1A and S1H). As with Mn^2+^, the optimal concentration of Co^2+^ and Ni^2+^ for the activation of nsp15 was approximately 5 mM (Figure 1A). The reactions with the nsp15 active-site mutant, H234A, demonstrated no cleavage of RNA substrates, indicating that the observed RNA degradation was not attributable to potential nuclease contamination or the ions and reagents themselves (Figure 1A). A 15 nt ssRNA substrate (ssRNA15) containing a single U site was employed in order to quantify the activation of nsp15 by Mn²⁺, Co²⁺ and Ni²⁺. Initially, three ion concentrations (0.05, 0.5 and 5 mM) were tested. However, due to the excessive activation of nsp15 by Co^2+^ and Ni^2+^, the method employed was unable to accurately quantify the activation effects of Co^2+^ and Ni^2+^ at 5 mM. Under identical conditions, the presence of 0.05 mM and 0.5 mM Mn^2+^ did not elicit a notable increase in nsp15 activity (Figure 1B). Conversely, the addition of 0.05 mM and 0.5 mM Co^2+^ resulted in a threefold and 17-fold enhancement of nsp15 activity, respectively, while the introduction of 0.05 mM and 0.5 mM Ni^2+^ led to an eightfold and 23-fold increase in nsp15 activity (Figure 1B). Moreover, the results of nsp15 cleavage of these three RNA substrates demonstrated that, to achieve an equivalent increase in nsp15 activity, Mn^2+^ was employed at a concentration approximately 30 times that of Co^2+^ and 100 times that of Ni^2+^ (Figure 1A and 1B). To further demonstrate the robust activation of nsp15 by 5 mM Co^2+^ and Ni^2+^, we utilized a fluorescence resonance energy transfer (FRET) assay. The cleavage of the FRET-RNA substrate by nsp15 would result in an increase in fluorescence. Different concentrations of nsp15 were employed in 5 mM EDTA, Mn^2+^, Co^2+^ and Ni^2+^ conditions to achieve a similar initial velocity (v_0_) of the cleavage reaction. The use of nsp15 under 5 mM EDTA was observed to be at least 75 times higher than under 5 mM Ni^2+^, 50 times higher than under 5 mM Co^2+^, and 7.5 times higher than under 5 mM Mn^2+^ (Figure 1C and S1I).

**Figure 1.**
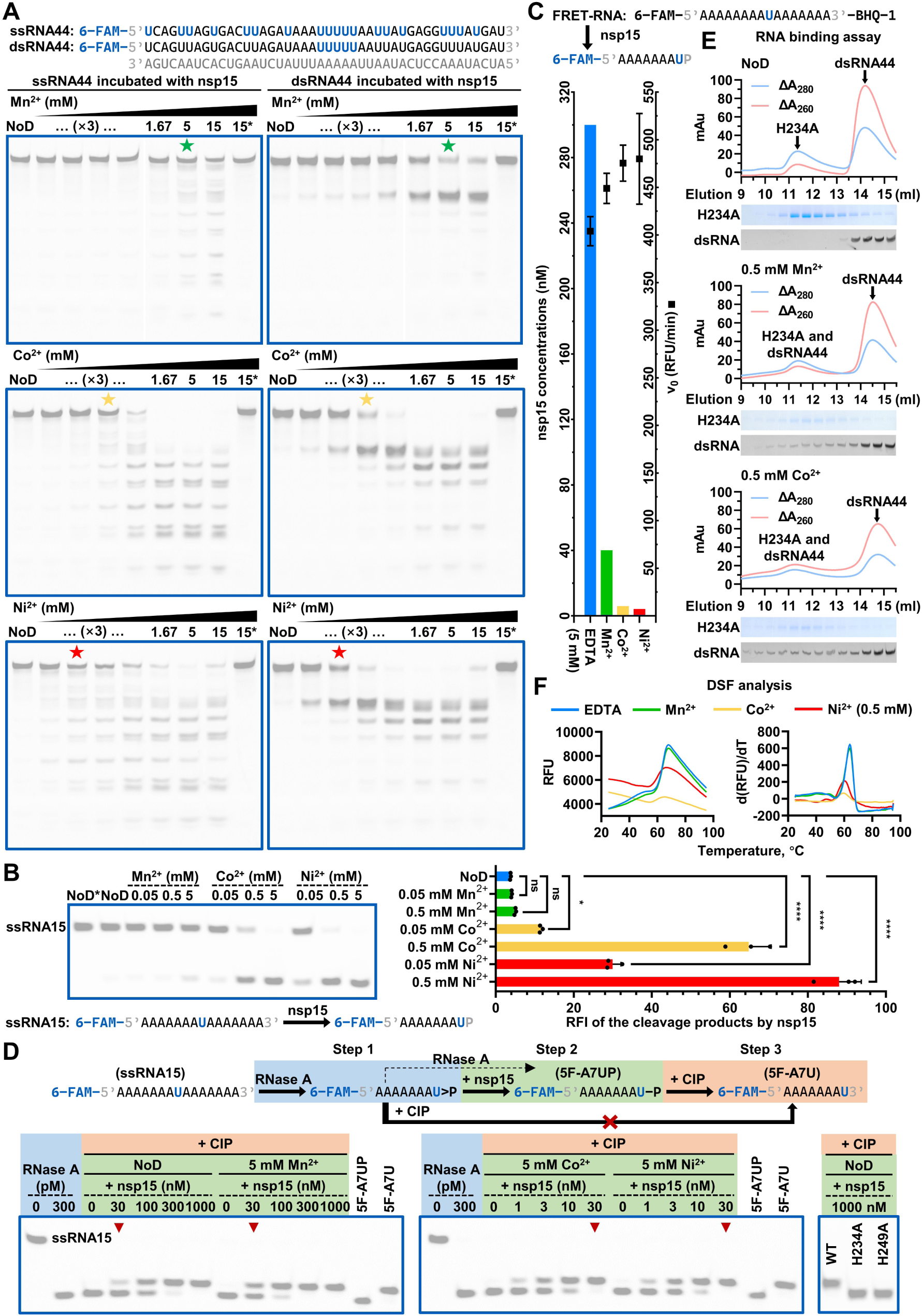
Co^2+^ and Ni^2+^ strongly activate SARS-CoV-2 nsp15. (A) Cleavage of ssRNA44 and dsRNA44 by 2 nM nsp15 in the presence of various concentrations of Mn^2+^, Co^2+^, or Ni^2+^. NoD refers to no divalent metal ions. The reactions containing the nsp15 active-site mutant, H234A, were used as negative controls, and these are marked with an asterisk (*). The sequences of ssRNA44 and dsRNA44 are displayed, with the cleavage sites preferred by nsp15 indicated in blue. The samples marked with pentagrams were used to analyze the preferred cleavage sites of nsp15 on ssRNA44 and dsRNA44 (see Figure S1H). (B) Quantification of the enhancing effects of Mn²⁺, Co²⁺, and Ni²⁺ on nsp15 EndoU activity using a single-U-containing substrate, ssRNA15. The ratio of the gray value of the cleavage product band to the sum of the gray values of the remaining substrate band and the cleavage product band is defined as the relative fluorescence intensity (RFI) of the cleavage products. The average and standard deviation of three independent reactions are graphed. Student’s t-test was performed. ns, not significant, *p* > 0.05; \**p* < 0.05; \*\*\*\**p* < 0.0001. (C) Quantification of the enhancing effects of Mn²⁺, Co²⁺, and Ni²⁺ on nsp15 EndoU activity using a FRET assay. The activity-enhancing effect is reflected by a decrease in the nsp15 concentration required to achieve a similar initial velocity (v_0_). The average and standard deviation of three independent reactions are graphed. (D) Hydrolysis of 2’,3’-cyclic phosphodiesters by nsp15. The reactions containing 30 nM nsp15 under different conditions are marked with red inverted triangles for easy comparison, suggesting the facilitation of the hydrolysis by Mn^2+^, Co^2+^, and Ni^2+^. (E) Enhanced RNA binding ability of nsp15 by Mn^2+^ and Co^2+^. The RNA binding assay is based on gel filtration chromatography. (F) Modified nsp15 stability by Co^2+^ and Ni^2+^. In the DSF analysis, each sample produced a melt curve (as shown on the left), and the first derivative of the melt curve produced a peak (as shown on the right), which provides the melting temperature (*T*_m_). RFU refers to relative fluorescence unit. See also Figure S1.

The initial study on SARS-CoV nsp15 reported that the 5’ products of RNA cleavage by nsp15 contained 2’,3’-cyclic phosphate ends.^24^ The subsequent study demonstrated that SARS-CoV nsp15 was capable of catalyzing both transphosphorylation of RNA to form a 2’,3’-cyclic phosphodiester and its subsequent hydrolysis to a 3’-phosphomonoester.^36^ A study of SARS-CoV-2 nsp15 proposed that 2’,3’-cyclic phosphate was the major cleavage product, with hydrolysis occurring at a slow rate.^35^ In order to investigate whether Co^2+^ and Ni^2+^ could also enhance the efficiency of nsp15 in hydrolyzing 2’,3’-cyclic phosphodiesters, we treated the cleavage products of ssRNA15 by nsp15 with calf intestine alkaline phosphatase (CIP), which can remove only 3’ phosphates.^24,36^ The end of the cleavage product can be identified as either a 2’,3’-cyclic phosphate or a 3’ phosphate by observing the change in electrophoretic mobility of the product due to the loss of a negative charge. It was unexpected that in the absence of any divalent metal ions, nsp15 exhibited high efficiency in the hydrolysis of 2’,3’-cyclic phosphodiesters, resulting in a relatively low accumulation of 2’,3’-cyclic phosphate ends (Figure S1J). We further demonstrated that DTT did not affect the hydrolysis reaction (Figure S1K). In the presence of Mn^2+^, Co^2+^, and Ni^2+^, both the transphosphorylation reaction and the hydrolysis reaction were correspondingly enhanced (Figure S1J). Consistently, the Co^2+^- and Ni^2+^-mediated activity enhancement was significantly stronger than that of Mn^2+^. To further investigate the enhancing effect of Co^2+^ and Ni^2+^ on the hydrolysis activity of nsp15 and to demonstrate that the hydrolysis reaction is a separate process, we utilized RNase A to prepare the 2’,3’-cyclic phosphate ends as substrates. RNase A is a well-characterized enzyme that has been proposed to catalyze both the transphosphorylation reaction and the hydrolysis reaction. However, the transphosphorylation rate is considerably faster than the hydrolysis rate, resulting in a notable accumulation of 2’,3’-cyclic phosphodiesters (Figure S1L).^37^ The use of the cleavage products with 2’,3’-cyclic phosphate ends by RNase A as substrates allows for a more precise observation of the pronounced enhancement of Co^2+^ and Ni^2+^ on the hydrolysis of 2’,3’-cyclic phosphodiesters by nsp15 (Figure 1D). In RNase A, the active-site residues H12 and H119 are responsible for the transphosphorylation reaction and the hydrolysis reaction via a general acid-base catalysis mechanism.^37^ Similarly, in nsp15, it has been proposed that the active-site residues H234 and H249 are the crucial determinants in the transphosphorylation reaction. We demonstrated that the H234 and H249 residues of nsp15 also played a pivotal role in the hydrolysis reaction (Figure 1D). This provides further evidence that nsp15 employs an RNase A-like catalytic mechanism.

Previous attempts to detect RNA binding to nsp15 using the electrophoretic mobility shift assay (EMSA) had not been successful.^17^ Consequently, in order to investigate the impact of divalent metal ions on the RNA binding ability of nsp15, gel filtration chromatography was employed, with the nsp15 active-site mutant, H234A, and dsRNA44. In the absence of divalent metal ions, no discernible interaction was observed between H234A and dsRNA44, and H234A and dsRNA44 were eluted separately, with H234A eluting early and dsRNA44 eluting late (Figure 1E and S1M). The presence of 0.5 mM Mn²⁺ resulted in the elution of some dsRNA44 in conjunction with H234A. This phenomenon was more pronounced in the presence of 0.5 mM Co²⁺ (Figure 1E). These results suggested that both Mn^2+^ and Co^2+^ (with Co^2+^ exhibiting superior efficacy) were capable of enhancing the interaction between the nsp15 protein and RNA substrates. However, attempts to detect the effect of Ni^2+^ on the RNA binding ability of nsp15 by the same method failed because high concentrations of nsp15 protein tended to precipitate at 0.5 mM Ni^2+^.

Our previous research had demonstrated that nsp15 stability remained consistent across different Mn^2+^ concentrations, including an EDTA control.^17^ We investigated the impact of Co^2+^ and Ni^2+^ on nsp15 stability by DSF. In contrast to the results observed with Mn^2+^, the melt curves of nsp15 in the presence of Co^2+^ and Ni^2+^ exhibited a markedly different profile compared to that observed in the presence of EDTA (Figure 1F). The first derivative of the melt curve produced a peak, which corresponded to the melting temperature (*T*_m_). The results indicated that the *T*_m_ of nsp15 in the presence of Co^2+^ and Ni^2+^ were lower than that in the presence of EDTA and Mn^2+^, suggesting that Co^2+^ and Ni^2+^ appeared to affect nsp15 structure and reduce its stability (Figure 1F).

### Zn^2+^ markedly inhibits both activities of SARS-CoV-2 nsp15 while Cu^2+^ irreversibly denatures nsp15 protein

In the initial examination of the impact of Zn^2+^ and Cu^2+^ on nsp15, it was observed that both metal ions markedly impeded the RNA cleavage activity of nsp15 (Figure 2A). It is noteworthy that the inhibitory effect of Zn^2+^ on nsp15 cleavage of dsRNA44 was more pronounced than that observed on nsp15 cleavage of ssRNA44 (Figure 2A). The inhibitory effect was quantified using the ssRNA15 substrate, which indicated that the presence of 0.05 mM and 0.5 mM Zn^2+^ resulted in a threefold and sevenfold reduction of nsp15 activity, respectively, while the introduction of 0.05 mM and 0.5 mM Cu^2+^ led to the complete elimination of nsp15 activity (Figure 2B). Furthermore, we demonstrated that Zn^2+^ and Cu^2+^ also exhibited a pronounced inhibitory effect on the activity of nsp15 in the hydrolysis of 2’,3’-cyclic phosphodiesters (Figure 2C). In order to ascertain whether the inhibition exerted by Zn^2+^ and Cu^2+^ is reversible, we initially incubated nsp15 with Zn^2+^ and Cu^2+^, subsequently removing these ions by the addition of excess EDTA, and then proceeded to test the transphosphorylation and hydrolysis activities of nsp15. The results indicated that following the removal of Zn^2+^, both activities of nsp15 were recovered. Nevertheless, nsp15 treated with Cu^2+^ did not display any observable activity after Cu^2+^ removal (Figure 2B and 2C). The DSF results showed that the melt curves of nsp15 in the presence of Zn^2+^ and Cu^2+^ were significantly different from that observed in the presence of EDTA (Figure 2D). As with Co^2+^ and Ni^2+^, the *T*_m_ of nsp15 in the presence of Zn^2+^ was lower than that in the presence of EDTA, suggesting the reduction of nsp15 stability by Zn^2+^ (Figure 2D). However, the first derivative of the melt curve of nsp15 in the presence of Cu^2+^ did not yield a pronounced peak (Figure 2D). Considering the irreversible elimination of nsp15 activity by Cu²⁺, we postulated that Cu²⁺ may induce nsp15 denaturation. We analyzed the nsp15 protein treated with Cu^2+^ and other divalent metal ions by Native-PAGE, the gel bands exhibited notable differences when comparing the protein treated with Cu^2+^ to those treated with other ions or in the absence of ions. The gel bands of the nsp15 protein treated with Cu^2+^ displayed an unusual ladder-like shape configuration, while the prominent bands of the nsp15 protein treated with other ions or in the absence of ions were slightly higher than the 228 kDa marker, consistent with the size of the natural hexameric form of nsp15 (Figure 2E and S2A). The SDS-PAGE analysis showed that the gel bands of the nsp15 protein, treated with Cu^2+^ or without ions, exhibited no discernible difference (Figure S2B). The findings suggested that Cu^2+^ altered the spatial configuration of nsp15 in an irreversible manner, with minimal impact on its primary structure. It is interesting to note that, while Zn^2+^ did not denature nsp15 protein, the presence of Zn^2+^ caused protein aggregation at a high nsp15 concentration (Figure 2F and S2C). A less pronounced protein aggregation phenomenon was observed in the presence of Co^2+^ and Ni^2+^ (Figure 2F and S2C), whereas this phenomenon was not observed in the presence of Mn^2+^ and in the absence of divalent metal ions (Figure 2F). This is consistent with previous observations in DSF experiments in which Co^2+^, Ni^2+^ and Zn^2+^ could reduce the stability of nsp15. The aggregation of nsp15 protein was eliminated when Co^2+^, Ni^2+^, and Zn^2+^ were removed by further addition of EDTA (Figure 2F), suggesting that the effects caused by these divalent metal ions were reversible. Given the observation that Zn^2+^ exerted the most significant effect on nsp15 stability in comparison to Co^2+^ and Ni^2+^, it was hypothesized that the specific interaction of Zn^2+^ with nsp15 might be the most potent. We examined the activation of nsp15 by Co^2+^ and Ni^2+^ in the presence of Zn^2+^. The results demonstrated that the addition of 0.05 mM Co^2+^ or Ni^2+^ under 0.05 mM Zn^2+^ did not result in an increase in RNA cleavage activity of nsp15 (Figure 2G). This finding suggested that the activating effect of Co^2+^ and Ni^2+^ was entirely counteracted by the inhibitory effect of Zn^2+^. In fact, only a slight increase in the RNA cleavage activity of nsp15 was observed with the addition of 0.5 mM Ni^2+^ in the presence of 0.05 mM Zn^2+^, whereas no such increase was observed with the addition of 0.5 mM Co^2+^ (Figure 2G). In addition, no increase in the RNA cleavage activity of nsp15 was observed upon the addition of 5 mM Mn^2+^ under 0.05 mM Zn^2+^ condition (Figure 2G). Furthermore, the addition of 5 mM Ni^2+^, Co^2+^ and Mn^2+^ at 0.05 mM Cu^2+^ did not result in any RNA cleavage by nsp15 (Figure S2D), indicating that Ni^2+^, Co^2+^ and Mn^2+^ did not influence the denaturation of nsp15 protein induced by Cu^2+^.

**Figure 2.**
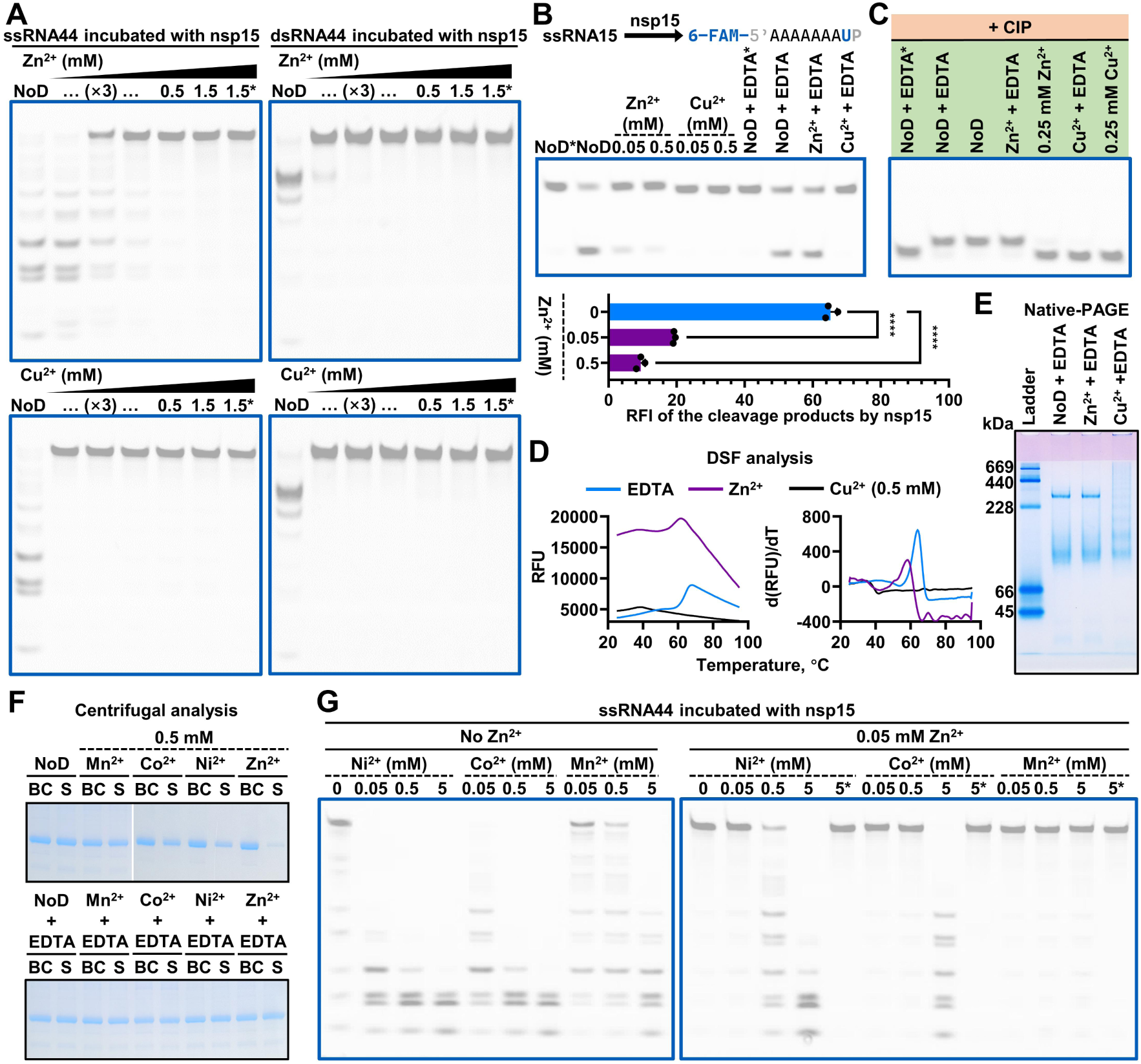
Zn^2+^ markedly inhibits both activities of SARS-CoV-2 nsp15 while Cu^2+^ irreversibly denatures nsp15 protein. (A) Cleavage of ssRNA44 and dsRNA44 by 200 nM nsp15 in the presence of various concentrations of Zn^2+^ or Cu^2+^. (B) Cleavage of ssRNA15 by nsp15 in the presence of Zn^2+^ or Cu^2+^, or by nsp15 pretreated with Zn^2+^ or Cu^2+^. In the former case, the average and standard deviation of three independent reactions are graphed. Student’s t-test was performed. ****p < 0.0001. In the latter case, 0.5 mM Zn^2+^ or Cu^2+^ was removed with 5 mM EDTA prior to the addition of ssRNA15. (C) Hydrolysis of 2’,3’-cyclic phosphodiesters by nsp15 in the presence of Zn^2+^ or Cu^2+^, or by nsp15 pretreated with Zn^2+^ or Cu^2+^. In the latter case, 0.5 mM Zn^2+^ or Cu^2+^ was removed with 5 mM EDTA prior to the addition of the products containing 2’,3’-cyclic phosphodiesters produced by the cleavage of ssRNA15 by RNase A. (D) Modified nsp15 stability by Zn^2+^ and Cu^2+^. (E) Native-PAGE analysis of the nsp15 protein pretreated with Zn^2+^ or Cu^2+^. The nsp15 hexamer has an estimated molecular weight of approximately 240 kDa. (F) Centrifugal analysis of the nsp15 sample with the presence of Mn^2+^, Co^2+^, Ni^2+^, or Zn^2+^, or with the removal of these ions through the subsequent addition of EDTA. In the latter case, 0.5 mM Mn^2+^, Co^2+^, Ni^2+^, or Zn^2+^ was removed with 5 mM EDTA prior to centrifugation. BC and S refer to before centrifugation and supernatant, respectively. (G) Cleavage of ssRNA44 by 20 nM nsp15 in the presence of Ni^2+^, Co^2+^, or Mn^2+^, with or without Zn^2+^. See also Figure S2.

The affinity of transition metal ions (Zn²⁺, Cu²⁺, Co²⁺, and Ni²⁺, among others) for specific amino acids (histidine and cysteine) has been documented since 1948.^38^ To eliminate the possibility that the specific effects of Zn²⁺, Cu²⁺, Co²⁺, and Ni²⁺ on nsp15 were due to interactions with the 6× His tag at the N terminus of the recombinant nsp15 protein, a Strep-Tag II was introduced in place of the 6× His tag to obtain the high-purity recombinant nsp15 protein without a 6× His tag and its active-site mutant (Figure S2E). The impact of Zn²⁺, Cu²⁺, Co²⁺, and Ni²⁺ on this Strep-nsp15 was tested. It was observed that Zn²⁺ and Cu²⁺ suppressed the RNA cleavage activity of nsp15, while Co²⁺ and Ni²⁺ enhanced it, as previously observed (Figure S2F). Given the similarity between nsp15 and RNase A in terms of the relative positions of the key active-site residues as well as the catalytic mechanism,^25,32,35,39,40^ we investigated whether Zn²⁺, Cu²⁺, Co²⁺, and Ni²⁺ exert a similar effect on RNase A as nsp15. The results indicated that Co^2+^ and Ni^2+^ did not enhance, but rather slightly inhibited the activity of RNase A (Figure S2G). Cu^2+^ exerted no discernible influence on RNase A, whereas Zn^2+^ demonstrated a modest inhibitory effect on the activity of RNase A, which was significantly weaker than that observed with nsp15 (Figure S2G).

### Specific effects of Co^2+^, Ni^2+^ and Zn^2+^ on nsp15 are widespread in coronaviruses

In addition to SARS-CoV-2, SARS-CoV, MERS-CoV, MHV-A59, porcine epidemic diarrhea virus (PEDV), and human coronavirus (HCoV)-229E are also commonly used in the study of coronaviruses. We purified the recombinant nsp15 proteins of these coronaviruses (Figure S3A) and investigated the impact of Mn^2+^, Co^2+^, Ni^2+^, Zn^2+^, and Cu^2+^ on them using ssRNA44 as substrates. It was demonstrated that in the absence of divalent metal ions, all of these nsp15 exhibited RNA cleavage activity (Figure 3A and S3B), indicating that this activity was not dependent on divalent metal ions. In the presence of 0.5 mM Zn^2+^, the RNA cleavage activity of all nsp15 was markedly suppressed, as observed in SARS-CoV-2 nsp15 (Figure 3A and S3B). Following the removal of Zn²⁺ by the addition of EDTA, the RNA cleavage activities of SARS-CoV nsp15, MHV-A59 nsp15 and HCoV-229E nsp15 were fully restored, whereas there was still a partial loss of RNA cleavage activity of MERS-CoV nsp15 and PEDV nsp15 (Figure S3C). Furthermore, the RNA cleavage activity of the majority of nsp15, excluding MHV-A59 nsp15, was irreversibly eliminated by Cu^2+^ (Figure 3A, S3B, and S3C), which was likely due to the potential denaturing effect of Cu^2+^ on nsp15 proteins. It was observed that the addition of 5 mM Co²⁺ or Ni²⁺ resulted in an enhancement of RNA cleavage activity in all nsp15 (Figure S3D). However, the addition of 5 mM Mn^2+^ did not result in a significant increase in the RNA cleavage activity of HCoV-229E nsp15 and PEDV nsp15 (Figure S3D). Given that both PEDV and HCoV-229E are classified as alphacoronaviruses, while other species are categorized as betacoronaviruses, it may be postulated that the activation of nsp15 by Mn^2+^ at the millimolar level was predominantly present in the latter group. The impact of 0.5 mM Mn²⁺, Co²⁺ and Ni²⁺ on nsp15 was quantified using the ssRNA15 substrate (Figure 3B). As observed in SARS-CoV-2 nsp15, the activation of SARS-CoV nsp15 and MERS-CoV nsp15 by 0.5 mM Mn^2+^ was not significant, whereas 0.5 mM Co^2+^ and Ni^2+^ demonstrated robust activation. The addition of 0.5 mM Co^2+^ and Ni^2+^ resulted in a 27-fold and 31-fold increase in SARS-CoV nsp15 activity, respectively, and a 22-fold and 26-fold increase in MERS-CoV nsp15 activity. In MHV-A59, the activation of nsp15 by Co^2+^ and Ni^2+^ was observed to be less pronounced than that observed for the other three betacoronaviruses. The addition of 0.5 mM Co^2+^ and Ni^2+^ resulted in an eightfold and 13-fold increase in MHV-A59 nsp15 activity, respectively. In contrast, the presence of 0.5 mM Mn^2+^ had a negligible impact on MHV-A59 nsp15. The activation of PEDV nsp15 and HCoV-229E nsp15 by Co^2+^ and Ni^2+^ was even weaker, with 0.5 mM Co^2+^ and Ni^2+^ demonstrating a fivefold and 12-fold activation in PEDV nsp15, respectively, and a fivefold and eightfold activation in HCoV-229E nsp15. Conversely, 0.5 mM Mn^2+^ exhibited no activation in either PEDV nsp15 or HCoV-229E nsp15. Moreover, 0.5 mM Mn^2+^ demonstrated an inhibitory effect on PEDV nsp15, although this inhibition was not statistically significant.

**Figure 3.**
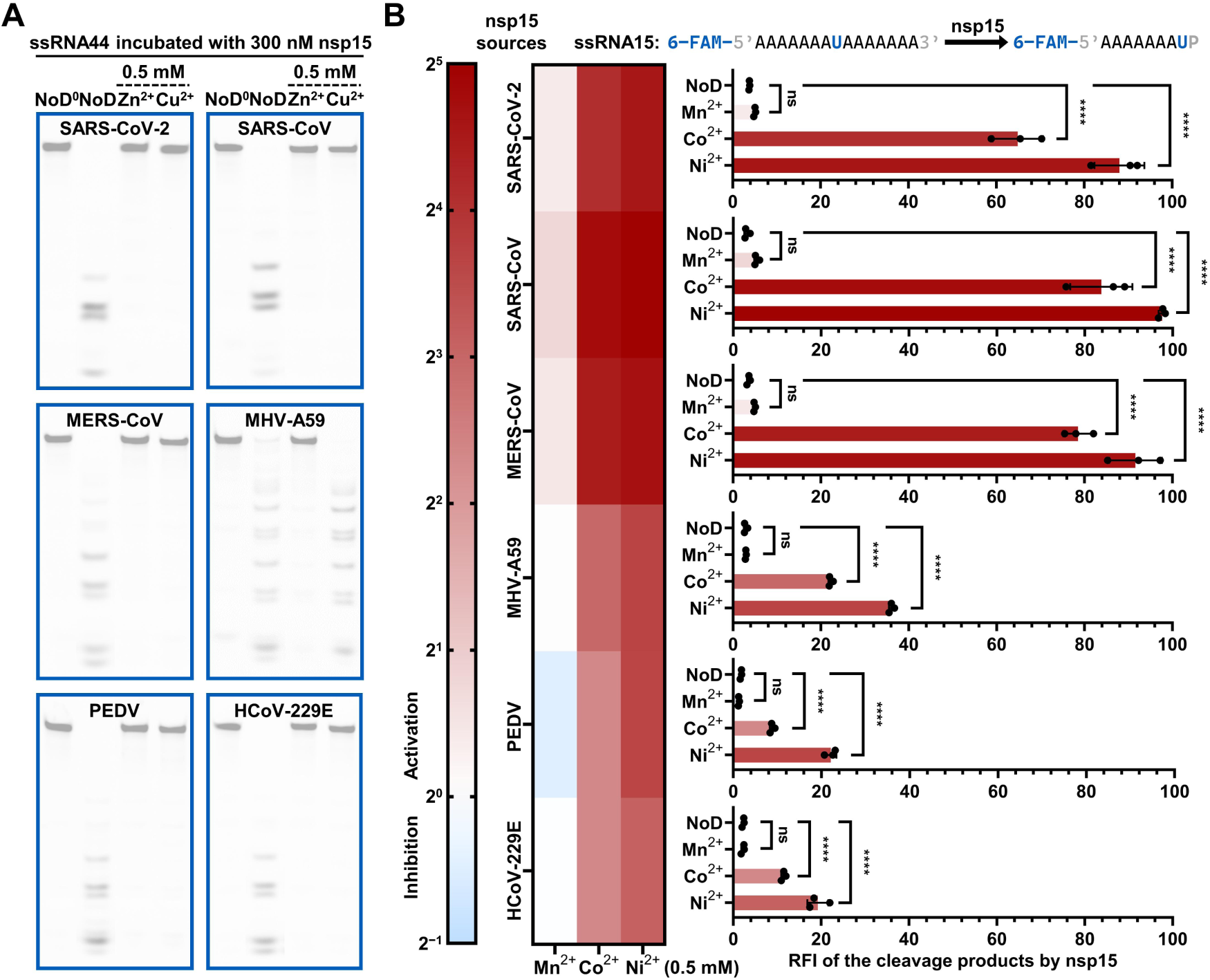
Specific effects of Co^2+^, Ni^2+^ and Zn^2+^ on nsp15 are widespread in coronaviruses. (A) Cleavage of ssRNA44 by nsp15 of various coronaviruses in the presence of Zn^2+^ or Cu^2+^. The reactions lacking nsp15 were used as negative controls, and these are marked with ^0^. (B) Quantification of the effects of Mn²⁺, Co²⁺, and Ni²⁺ on the EndoU activity of nsp15 of various coronaviruses, using a single-U-containing substrate, ssRNA15. The average and standard deviation of three independent reactions are graphed (as shown on the right). Student’s t-test was performed. ns, not significant, p > 0.05; ****p < 0.0001. The fold increase or decrease in nsp15 EndoU activity in the presence of Mn^2+^, Co^2+^, or Ni^2+^ is shown by a heatmap (as shown on the left). See also Figure S3.

### Co^2+^ enhances binding between SARS-CoV-2 nsp15 and dsRNA

As previously stated, no divalent metal ions or metal-binding sites had been identified in any of the nsp15 structural studies, with the presence of Mn^2+^ in the majority of them.^26,34,35^ Given the significant effects of Co^2+^, Ni^2+^ and Zn^2+^ on the activities of SARS-CoV-2 nsp15, we attempted to capture the SARS-CoV-2 nsp15 structures in complex with dsRNA substrates in the presence of Co^2+^, Ni^2+^ or Zn^2+^ using cryo-EM, aiming to determine the metal-binding sites of nsp15 and to elucidate the regulatory mechanism of nsp15 activities by divalent metal ions. In order to prevent dsRNA cleavage and potential interactions between these three divalent metal ions and the 6× His tag, we purified SARS-CoV-2 nsp15 with H234A mutation and a N-terminal Strep-Tag II instead of a 6× His tag (referred to as Strep-H234A hereafter). We incubated Strep-H234A with dsRNA44 substrate in the presence of Co^2+^, Ni^2+^ or Zn^2+^. Both Zn^2+^ and Ni^2+^ induce protein aggregation, consistent with the previous results (Figure 2F), thus preventing the continuation of the structural studies with Zn^2+^ and Ni^2+^. We carried out cryo-EM imaging on the nsp15/dsRNA sample with Co^2+^. Notably, unlike previously reported cryo-EM 2D class averages that predominantly showed only one dsRNA bound per nsp15 hexamer,^41,42^ the 2D class averages of the nsp15/dsRNA in the presence of Co^2+^ revealed different nsp15/dsRNA complexes with up to three dsRNAs bound to one nsp15 hexamer (Figure 4A). This suggested that the presence of Co^2+^ enhanced the binding between nsp15 and dsRNA, which is consistent with our previous results (Figure 1E). Among all nsp15/dsRNA complexes, we successfully obtained high-resolution structures for the complexes containing one and two dsRNA molecules (6:1 and 6:2 nsp15/dsRNA complexes; Figure S4A–S4E). The overall resolution of the 6:1 nsp15/dsRNA complex is 2.6 Å, while the 6:2 complex has a resolution of 2.78 Å. In both complexes, nsp15 forms a hexamer composed of two back-to-back trimers with clear densities of bound dsRNAs (Figure 4B–4E). Each dsRNA engages three of the six nsp15 protomers, forming two distal interfaces and the major interface at the cleavage site of the dsRNA (Figure 4F). The first distal interface (Interface I) is mediated by one of the top nsp15 protomers and involves K12 and Q18 in the middle domain (MD) as well as S147 in the N-terminal domain (NTD) (Figure 4G). The second distal interface (Interface II) is mediated by one of the bottom nsp15 protomers and involves K110, T112, R135 and N136 in the MD (Figure 4H). The interactions at the major interface (Interface III) are mediated by the catalytic EndoU domain of another top nsp15 protomer, and the residues that interact with dsRNA are analogous to those previously reported, including H242, S243, H249, K289, S293, S315, W332, K334, E339, Y342, and K344 (Figure 4I).^41,42^ Furthermore, we observed a uridine (U18) that flips out from the dsRNA helix and adopts a position in the active site, similar to what has been observed in the UMP-bound nsp15 structure and the dsRNA-bound nsp15 structure (Figure 4J and 4K).^26,41^

**Figure 4.**
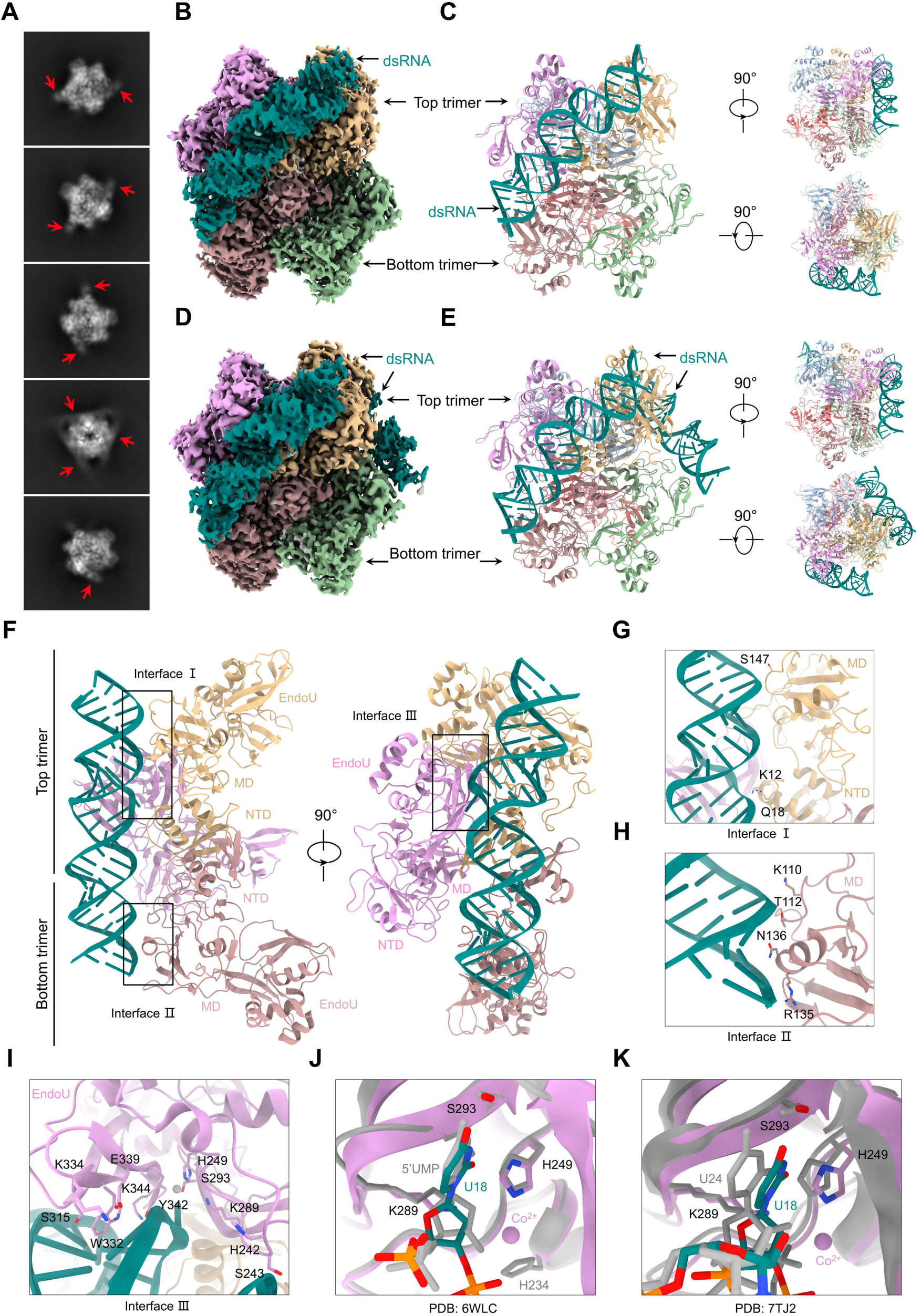
Structures of dsRNA-bound SARS-CoV-2 nsp15 in the presence of Co^2+^. (A) 2D class averages of different nsp15/dsRNA complexes. dsRNA is indicated by red arrows. (B) Cryo-EM density map of the 6:1 nsp15/dsRNA complex. (C) Ribbon diagram of the 6:1 nsp15/dsRNA complex in three orthogonal views. (D) Cryo-EM density map of the 6:2 nsp15/dsRNA complex. (E) Ribbon diagram of the 6:2 nsp15/dsRNA complex in three orthogonal views. (F) Ribbon diagram of the dsRNA binding interfaces in two orthogonal views. The dsRNA engages three of the six nsp15 protomers. (G) Enlarged view of the first distal interface formed by the NTD and MD domains of nsp15 (burly wood). (H) Enlarged view of the second distal interface formed by the MD domain of nsp15 (rosy brown). (I) Enlarged view of the major interface formed by the EndoU domain of nsp15 (plum). (J) Enlarged view of the overlaid ribbon diagram of the UMP-bound nsp15 (PDB 6WLC; gray) and our dsRNA-bound nsp15 (colored). (K) Enlarged view of the overlaid ribbon diagram of another dsRNA-bound nsp15 (PDB 7TJ2; gray) and our dsRNA-bound nsp15 (colored). See also Figure S4.

### Co^2+^ locates at the nsp15 active site to interact with dsRNA

Although metal ions are required by nsp15 to catalyze the cleavage of RNA, no metal ion density has been observed at the nsp15 active site in previous studies. Intriguingly, we observed an ion density at the nsp15 active site and modeled it as a Co^2+^ as it is the only divalent metal ion in the cryo-EM sample. The Co^2+^ is coordinated by H249 and K289 in the active site as well as the 2’ hydroxyl of the flipped-out U18 (Figure 5A). By overlaying the UMP-bound nsp15 complex with our structure, we found that H234, which was mutated to be alanine in our structure, also should participate in Co^2+^ coordination (Figure 5B). Another residue T340 was identified in proximity to the Co^2+^ (Figure 5B). Interestingly, the structural alignment of the previously reported nsp15/dsRNA complex and our Co^2+^-bound nsp15/dsRNA complex revealed an inward shift of the phosphate between the flipped-out U18 and A19 upon Co^2+^ binding (Figure 5C). This conformational change is necessary for ion binding at the active site or otherwise the 3’ phosphate of the U18 would clash with the ion. To demonstrate the interactions between the residues and the Co^2+^ in the active site, we created the H249A, K289A and T340A mutants (Figure S5A) and detected the RNA cleavage activity of these nsp15 mutants against dsRNA44 in the presence of Co^2+^. As anticipated, mutation of H249 and K289, the pivotal amino acids that facilitate the RNA cleavage reaction, significantly diminished the ability of nsp15 to cleave dsRNA44 (Figure 5D). Unexpectedly, mutation of T340, which is not the key residues, also led to a substantial decrease in nsp15 activity (Figure 5D). We hypothesized that T340A alters the local configuration of the active site due to proximity. The attempt to quantify the activation of K289A induced by Co^2+^ was unsuccessful due to the RNA cleavage by K289A in the absence of Co²⁺ being too weak to detect, as with H234A (Figure 5D). However, we found that the activation of H249A by Co^2+^ was considerably less pronounced in comparison to the wild-type (WT) nsp15 (Figure 5D). Conversely, T340A does not change the effect of Co²⁺ (Figure 5D). These findings demonstrate that Co^2+^ binds to nsp15’s active site to activate nsp15.

**Figure 5.**
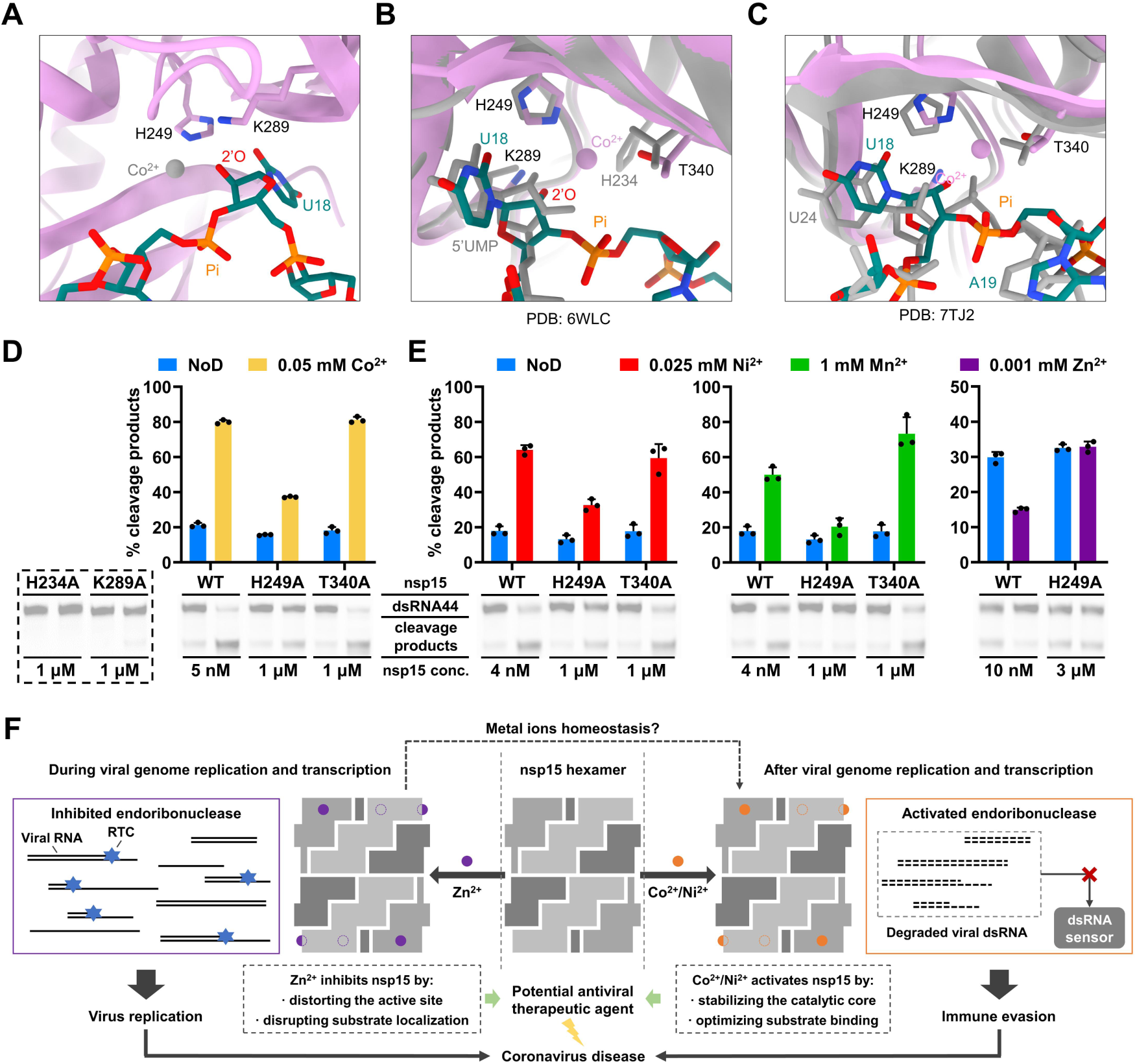
Interactions between divalent cations and nsp15. (A) Enlarged view of the catalytic residues at the nsp15 active site (plum) and the flipped-out U18 at the bound-dsRNA (teal) which can interact with Co^2+^ (gray). (B) Enlarged overlaid ribbon diagram of the Co^2+^ binding interface between the UMP-bound nsp15 complex (PDB 6WLC; gray) and our Co^2+^-bound nsp15/dsRNA complex (colored). (C) Enlarged view of the overlaid ribbon diagram, showing the conformational changes of the previously reported nsp15/dsRNA complex (PDB 7TJ2; gray) to our Co^2+^-bound nsp15/dsRNA complex (colored). (D, E) Impact of mutations in residues near the bound-Co^2+^, including H234, H249, K289, and T340, on the functions of divalent metal ions. The ratio of the gray value of the cleavage product band to the sum of the gray values of the remaining substrate band and the cleavage product band is defined as % cleavage products. The average and standard deviation of three independent reactions are graphed. (F) Model depicting the mechanism and physiological significance of metal ions in regulating the activity of the coronavirus nsp15. See also Figure S5.

To rule out other possible Co^2+^ binding site, we tested three putative Co^2+^ binding sites other than the active site based on electron densities. One is located in proximity to D282, S261, and P262; another is situated in the vicinity of H14, R61, and I63; and a third is close to H242. We generated D282A, S261A, H242A, and H14A mutants (Figure S5A), and observed a range of attenuated activities in these mutants compared to wild-type nsp15, with D282A showing the most significant attenuation (Figure S5B and S5C). Nevertheless, the enhancing effects of Co^2+^ on the activities of these mutants were similar to that on the wild-type nsp15 activity (Figure S5B), with the exception of D282A, where the enhancing effect of Co^2+^ on its activity was somewhat decreased (Figure S5D). Interestingly, a previous structural study of SARS-CoV-2 nsp15 had also reported the electron density near D282, S262, and P262, which was modeled as a Mg^2+^.^33^ However, no corresponding mutants were created to validate the interactions in the work. It is noteworthy that the electron density in proximity to D282, S262, and P262 in our structure was only detected in a single protomer of the nsp15 hexamer. Additionally, while mutation of D282 exhibited a certain degree of influence on the impact of Co^2+^, mutation of S262 demonstrated minimal effect, suggesting that this electron density may corresponds to ions other than Co^2+^.

### Ni^2+^, Zn^2+^, and Mn^2+^ function by binding to nsp15 active site

Given the finding that Co^2+^ binds to the nsp15 active site, the subsequent investigation aimed to ascertain whether other divalent metal ions would demonstrate a similar behavior within the active site. To this end, we examined the effects of Ni^2+^, Mn^2+^, and Zn^2+^ on the activity of H249A and T340A. Similar to Co^2+^, the enhancing effect of Ni²⁺ on H249A activity was less pronounced than its effect on wild-type nsp15 activity, whereas the effect of Ni^2+^ on T340A activity was comparable to that on wild-type nsp15 activity (Figure 5E). The enhancing effect of Mn^2+^ on H249A activity was also attenuated compared to wild-type nsp15. However, mutation of T340 significantly strengthened the enhancing effect of Mn^2+^ on nsp15 activity, an outcome that was not anticipated (Figure 5E). Given the fact that the ionic radius of Mn^2+^ is the most substantial among the three ions, it was hypothesized that, despite the non-directive involvement of T340 in the coordination of divalent metal ions, it may function as a spatial hindrance, thereby reducing the likelihood of Mn^2+^ binding to the active site in comparison to Co^2+^ and Ni^2+^. Consequently, when the threonine is mutated to an alanine, the side chain of the residue is shorter, thereby facilitating the binding of Mn^2+^ to the active site and its subsequent role. Notably, mutation of H249 led to a reduction in the inhibitory effect of Zn^2+^ on nsp15 activity (Figure 5E). This suggests the possibility that Zn^2+^ may also function by binding to the active site. It was found that the activating effect of Co^2+^ or Ni^2+^ on nsp15 could be completely counteracted by the inhibitory effect of Zn^2+^ when the two ions coexist at the same concentration (Figure 2G). We hypothesized that this phenomenon is related to the binding affinities of the metal ions to the active site.

Notably, we employed AlphaFold3 to predict the structures of complexes of divalent metal ions and nsp15. The predicted overall structure of nsp15 is similar to the actual structure (Figure S5E). Intriguingly, the predicted structure of Mn^2+^-bound nsp15 suggests that Mn^2+^ is localized in the active site, in close proximity to H234, H249, and K289 (Figure S5E and S5F). This observation is analogous to our cryo-EM structure. Conversely, the predicted Co^2+^-bound nsp15 structure indicates that Co^2+^ binds at a distinct site, situated in proximity to the interface between the two nsp15 trimers, near E56, C102, and S103 (Figure S5E and S5F). However, we demonstrated that the simultaneous mutation of E56, C102, and S103 (designated as mIII) did not modify the effect of Mn^2+^, Co^2+^, Ni^2+^, and Zn^2+^ on nsp15 activity (Figure 1B, 2B and S5G).

## DISCUSSION

It is generally accepted that nsp15 functions through a conventional acid-base catalysis mechanism analogous to that of RNase A. This assertion is supported by the observation that the active sites of these two enzymes are similar, with both comprising two histidine residues and a lysine residue. However, the differential regulation of nsp15 activity by transition metal ions—Co²⁺ and Ni²⁺ as activators and Zn²⁺ as an inhibitor—suggests a nuanced interplay between metal coordination chemistry and enzymatic catalysis in nsp15, which is not observed in RNase A. Specifically, Co²⁺ and Ni²⁺, with their six-coordinated octahedral coordination preferences, are likely to optimize the spatial arrangement of H234, H249, and K289 by coordinating with these three catalytic residues. In addition, the Co^2+^/Ni^2+^ present in the active site could interact with the 2’ hydroxyl of the positioned uridine and induce a local conformational change in RNA, increasing the likelihood of RNA hydrolysis. Conversely, the robust tetrahedral coordination geometry of Zn^2+^ potentially distort the nsp15 active site or disrupt the substrate pose to impede RNA hydrolysis.

The regulatory mechanism of nsp15 activity is physiologically crucial in coronavirus replication. As previously stated, it is essential to guarantee that viral RNA cannot be degraded by nsp15 during genome replication and transcription, while residual viral dsRNA must be efficiently degraded by nsp15 to avoid being detected by host cell dsRNA sensors following the completion of genome replication and transcription. Based on our findings, we hypothesized that local concentrations of metal ions in different cellular contexts govern nsp15’s activity. When nsp15 adopts a Zn^2+^-bound conformation, its endoribonuclease activity is suppressed to prevent degradation of viral RNA and allow viral genome replication and transcription. When nsp15 is in Co^2+^/Ni^2+^-bound form, viral dsRNA are efficiently removed to prevent host immune activation (Figure 5F). It is noteworthy that structural studies of SARS-CoV-2 nsp12, nsp13 and nsp14, which are an RNA-dependent RNA polymerase, a helicase and an exoribonuclease, respectively, and perform essential functions in viral genome replication and transcription, as well as nsp10, the cofactor for nsp14 and nsp16, have revealed that Zn^2+^ plays a structural role in maintaining the correct conformation of these nsps.^43–47^ These findings align with our hypothesis that nsp15 adopts a Zn^2+^-bound conformation during genome replication and transcription, in consideration of the co-localization of nsp15 and viral RTCs.^10,20–22^ However, there is a paucity of research on the effects of Co^2+^ and Ni^2+^ on other coronavirus nsps, which merits further investigation. In accordance with our hypothesis, interference with the regulation of nsp15 activity, whether its activation or inhibition, has the potential to affect viral genome replication and transcription, as well as nsp15-mediated immune evasion. Accordingly, the metal-binding site of nsp15 represents a promising target for antiviral drug design. Zn^2+^, Co^2+^ and Ni^2+^ or their regulators might be used to treat coronavirus disease as a broad-spectrum nsp15 inhibitor or activator. Actually, zinc has been proposed as a potential therapeutic agent for the treatment of COVID-19 due to its immunomodulatory and antiviral properties, and zinc therapy has been demonstrated to exhibit beneficial effects on patients infected with SARS-CoV-2.^48–51^ Our findings suggest that the effectiveness of zinc therapy is attributable not only to its role in mediating the host antiviral immune response, but also potentially to its direct action on viral proteins.

However, the underlying mechanisms that precipitate the shift of nsp15 metal-binding state remain to be elucidated. Intracellular zinc homeostasis may be a key clue in this regard. The total concentration of zinc within cells is estimated to range from 200 to 300 μM, with the majority being protein-bound.^52^ The concentration of free intracellular zinc is subject to stringent regulation, maintained within the range of high picomolar to low nanomolar levels.^53–55^ Zinc transporters (ZnTs) and Zrt/Irt-related proteins (ZIPs) facilitate zinc influx, efflux, and compartmentalization across biological membranes, thereby regulating zinc homeostasis within cells.^56,57^ Zinc homeostasis has been demonstrated to be crucial for the maintenance of innate immunity.^58–63^ We hypothesize that the distribution of intracellular Zn²⁺ and the changes in its concentration might play an important regulatory role in the replication process of coronaviruses. The roles of ZnTs and ZIPs in the viral replication process are worthy of exploration. Interestingly, a recent study revealed extensive alterations in metal homeostasis, including cobalt and nickel, in the body during the course of the SARS-CoV-2 infection.^64^ Accordingly, further study is necessary to determine the cobalt and nickel homeostasis during such an infection, in consideration of the specific impact of Co^2+^ and Ni^2+^ on nsp15.

## METHODS

### Protein expression and purification

#### For wild-type SARS-CoV-2 nsp15 and its mutants

The wild-type SARS-CoV-2 nsp15 (YP_009725310.1) with an N-terminal 6× His tag and its H234A mutant were expressed and purified as previously described (complete sequence of the nsp15 expression plasmid is listed in Table S1).^17^ A Strep-Tag II (Trp-Ser-His-Pro-Gln-Phe-Glu-Lys) was introduced in place of the 6× His tag using the Gibson assembly method to obtain the Strep-nsp15 and Strep-H234A constructs (primers for cloning are listed in Table S2). The constructs were transformed into *E. coli* BL21(DE3)pLysS cells (ToloBio, China). These cells were cultured in 1L (for Strep-H234A) or 4 L (for Strep-nsp15) of LB medium containing 50 µg/mL kanamycin and 34 µg/mL chloramphenicol at 37°C until an OD600 of approximately 0.8 was reached. The flasks containing the culture were then placed at 4°C for 1 h. Subsequently, protein expression was induced by addition of 0.2 mM IPTG, and incubation continued at 16°C for 16 h. The cells were harvested and then stored at −80°C. The frozen cells were thawed on ice and resuspended in a Strep binding buffer [100 mM Tris-HCl (pH 8.0 at 25°C), 300 mM NaCl, and 1 mM EDTA], then lysed by ultrasonication. The supernatant was collected after centrifugation at 14,000 rpm and 4°C for 1 h and filtered with 0.45-μm filters. The filtered supernatant was loaded onto a StrepTrap XT 1 mL column (Cytiva, USA) pre-equilibrated with 10 volumes of the Strep binding buffer. The column was washed with 50 volumes of the Strep binding buffer. Most of the Strep-tagged nsp15 was eluted by a Strep binding buffer that had been supplemented with 40 mM biotin. The collected eluates were concentrated with an Amicon Ultra-15 30K centrifugal filter (Millipore, USA) and dialyzed against a storage buffer containing 50 mM Tris-HCl (pH 8.0 at 25°C), 100 mM NaCl, 0.1% (v/v) Triton X-100, and 50% (v/v) glycerol, and then stored at −20°C.

Amino acid mutations (H14A, H242A, H249A, S261A, D282A, K289A, T340A and mIII) were introduced on the His-nsp15 construct using the Gibson assembly method (primers for cloning are listed in Table S2). Theses mutants were expressed with the same procedure as detailed above and purified using an Ni-NTA agarose column (Qiagen, Germany). Specifically, the frozen cells were thawed on ice and resuspended in a lysis buffer [50 mM Tris-HCl (pH 8.0 at 25°C), 500 mM NaCl, 5% (v/v) glycerol, and 10 mM imidazole], then lysed by ultrasonication. The supernatant was collected after centrifugation at 14,000 rpm and 4°C for 1 h and filtered with 0.45-μm filters. The filtered supernatant was loaded onto the Ni-NTA agarose column pre-equilibrated with five volumes of the lysis buffer. The column was first washed with 10 volumes of an elution buffer [20 mM Tris-HCl (pH 8.0 at 25°C), 500 mM NaCl, 5% (v/v) glycerol, and the indicated concentration of imidazole] containing 30 mM imidazole, and then five volumes of an elution buffer containing 50 mM, 60 mM, 70 mM, 80 mM, and 200 mM imidazole. The eluates with 70 mM, 80 mM, and 200 mM imidazole were collected and mixed, then concentrated and dialyzed as previously described.

#### For nsp15 of other coronaviruses

The DNA fragments encoding nsp15 of SARS-CoV (NP_828872.1), MERS-CoV (YP_009047226.1), MHV-A59 (YP_009915685.1), PEDV (NP_839968.1), and HCoV-229E (NP_835355.1) were synthesized and inserted into pET-28a(+) vectors harboring an N-terminal 6× His tag by GenScript (complete sequences of the nsp15 expression plasmids are listed in Table S1). SARS-CoV nsp15, MERS-CoV nsp15, MHV-A59 nsp15, and HCoV-229E nsp15 were expressed with the same procedure as detailed above, with the exception of the use of Rosetta 2(DE3)pLysS cells (TOLOBIO, China), and were purified using the Ni-NTA agarose column as previously described, excepting that only eluates with 80 mM and 200 mM imidazole were collected. The expression and purification process of PEDV nsp15 is somewhat different. First, the protein expression of PEDV nsp15 was induced by addition of 1 mM IPTG, and incubation continued at 25°C for 16 h. In addition, a second purification step was performed. Specifically, the eluates with 70 mM, 80 mM, and 200 mM imidazole were collected, followed by their mixing and concentration. Subsequently, the concentrated sample was loaded onto a Superdex 200 Increase 10/300 GL column (Cytiva, USA) pre-equilibrated with a gel filtration buffer [20 mM Tris–HCl (pH 8.0 at 25°C), 500 mM NaCl, and 5% (v/v) glycerol] for gel filtration chromatography.

Protein concentrations were determined with a Bradford Protein Quantitative kit (Bio-Rad, USA), with bovine serum albumin as a standard. Protein purity was analyzed by SDS-PAGE with Coomassie blue staining. Protein size was determined with PageRuler Plus Prestained Protein Ladder (Thermo Scientific, USA) in SDS-PAGE analysis.

#### RNA substrates preparation

The RNA oligonucleotides, including ssRNA15, ssRNA44, fluorescence resonance energy transfer (FRET)-RNA, and the complementary strand of ssRNA44, were synthesized chemically by GenScript. dsRNA44 was generated through the process of annealing between ssRNA44 and its complementary strand. The RNA annealing reaction containing 10 mM Tris-HCl (pH 7.4 at 25°C), 50 mM KCl, 25 μM ssRNA44, and 25 μM the complementary strand of ssRNA44, was carried out on a PCR instrument with a specific annealing program (1. 90°C, 1 min; 2. 90°C, 5 s, −0.1°C per cycle; 3. GOTO step 2, 650×; 4. 4°C, ∞.).

### RNA cleavage assays

#### ssRNA44 or dsRNA44 as substrates

500 nM ssRNA44 or dsRNA44 was incubated with the indicated concentration of nsp15 to a final volume of 10 μL in an RNA cleavage buffer [50 mM Tris-HCl (pH 7.4 at 25°C), 140 mM KCl, and indicated concentrations of MnCl_2_, CoCl_2_, NiCl_2_, ZnCl_2_, and CuCl_2_]. The reactions were performed at 37°C for the indicated time (30 min if not otherwise specified) and then terminated by the addition of 10 μL of 2× RNA loading dye (New England Biolabs, USA). The samples were heated at 85°C for 2 min, immediately placed on ice for 2 min. To generate an RNA size ladder, alkaline hydrolysis of ssRNA44 at a concentration of 6 μM to a final volume of 10 μL in an alkaline hydrolysis buffer [50 mM sodium carbonate (pH 9.4) and 1 mM EDTA] was performed for 15 min at 90°C and quenched with 10 μL of 2× RNA loading dye. All samples were analyzed by 20% TBE-urea PAGE (8 M urea). RNA was visualized with a ChemiScope imager (CLiNX, China) using the Cy2 (Ex470BL, Em525/30F) channel. Images were processed and the grey values of gel bands were quantified with ImageJ software.

To investigate the impact of the metal ions on RNase A activity, 500 nM ssRNA44 was incubated with 5 pM RNase A (Thermo Scientific, USA) to a final volume of 10 μL in an RNA cleavage buffer [50 mM Tris-HCl (pH 7.4 at 25°C), 140 mM KCl, and zero or 0.5 mM MnCl_2_, CoCl_2_, NiCl_2_, ZnCl_2_, or CuCl_2_]. The reactions were performed at 37°C for 30 min and then terminated by the addition of 10 μL of 2× RNA loading dye. The samples were heated at 85°C for 2 min, immediately placed on ice for 2 min, and then analyzed by 20% TBE-urea PAGE (8 M urea). RNA was visualized with a ChemiScope imager using the Cy2 (Ex470BL, Em525/30F) channel. Image processing was conducted using ImageJ software.

#### ssRNA15 as substrates

500 nM ssRNA15 was incubated with the indicated concentration of nsp15 to a final volume of 10 μL in an RNA cleavage buffer [50 mM Tris-HCl (pH 7.4 at 25°C), 140 mM KCl, and the indicated concentrations of MnCl_2_, CoCl_2_, NiCl_2_, ZnCl_2_, or CuCl_2_]. The reactions were performed at 37°C for 15 min and then terminated by the addition of 10 μL of 2× RNA loading dye. The samples were heated at 85°C for 2 min, immediately placed on ice for 2 min, and then analyzed by 20% TBE-urea PAGE (8 M urea). RNA was visualized with a ChemiScope imager using the Cy2 (Ex470BL, Em525/30F) channel. Images were processed and the grey values of gel bands were quantified with ImageJ software. The concentrations of SARS-CoV-2 nsp15, SARS-CoV nsp15, MERS-CoV nsp15, MHV-A59 nsp15, PEDV nsp15, and HCoV-229E nsp15 used in the quantification study of the enhancing effects of Mn^2+^, Co^2+^, and Ni^2+^ on nsp15 EndoU activity, were 20 nM, 30 nM, 175 nM, 225 nM, 20 nM, and 25nM, respectively. The concentration of SARS-CoV-2 nsp15 used in the quantification study of the inhibitory effect of Zn^2+^ on nsp15 EndoU activity was 1000 nM.

#### FRET-RNA as substrates

500 nM FRET-RNA was incubated with the indicated concentration of nsp15 to a final volume of 20 μL in an RNA cleavage buffer [50 mM Tris-HCl (pH 7.4 at 25°C), 140 mM KCl, and 5 mM EDTA, MnCl_2_, CoCl_2_, or NiCl_2_] at 37°C. The cleavage of the FRET-RNA substrate would result in an increase in fluorescence. Time-course reactions were directly monitored in a CFX Connect qPCR system (Bio-Rad, USA) using the FAM channel. The initial velocity (v_0_) was estimated using the slope from linear regressions of the reaction monitored during 1–5 min.

#### Detection of the hydrolysis of 2’,3’-cyclic phosphodiesters For RNase A

500 nM ssRNA15 was incubated with the indicated concentration of RNase A and 5U Quick CIP (calf intestine alkaline phosphatase, New England Biolabs, USA) to a final volume of 10 μL in an rCutSmart buffer (New England Biolabs, USA). The reactions were performed at 37°C for 30 min and then terminated by the addition of 10 μL of 2× RNA loading dye. The samples were heated at 85°C for 2 min, immediately placed on ice for 2 min.

#### For nsp15

**Method 1.** The RNA cleavage reaction with ssRNA15 as substrate (detailed above) was terminated by heating the sample at 85°C for 5 min. Subsequently, 1.25 μL of 10× rCutSmart buffer and 1.25 μL of Quick CIP (5U/μL) were added into the sample. The reactions were performed at 37°C for 30 min and then terminated by the addition of 12.5 μL of 2× RNA loading dye. The samples were heated at 85°C for 2 min, immediately placed on ice for 2 min.

**Method 2**. 500 nM ssRNA15 was incubated with 300 pM RNase A in an RNA cleavage buffer [50 mM Tris-HCl (pH 7.4 at 25°C) and 140 mM KCl] at 37°C for 15 min to produce 2’,3’-cyclic phosphodiesters. For the reaction with Mn^2+^, Co^2+^, or Ni^2+^, 8 μL of the RNase A reaction sample was added with zero or 5 mM MnCl_2_, CoCl_2_, or NiCl_2_, and the indicated concentration of nsp15, to a final volume of 10 μL; for the reaction with Zn^2+^ or Cu^2+^, 5 μL of the RNase A reaction sample was added with 5 μL of the mixture containing zero or 0.5 mM ZnCl_2_ or CuCl_2_, and 1000 nM nsp15. The reaction was then incubated at 37°C for 15 min. Subsequently, 1.25 μL of 10× rCutSmart buffer and 1.25 μL of Quick CIP (5U/μL) were added into the sample. The reactions were performed at 37°C for 30 min and then terminated by the addition of 12.5 μL of 2× RNA loading dye. The samples were heated at 85°C for 2 min, immediately placed on ice for 2 min.

All samples were analyzed by 20% TBE-urea PAGE (8 M urea) with 5F-A7UP and 5F-A7U as markers. RNA was visualized with a ChemiScope imager using the Cy2 (Ex470BL, Em525/30F) channel. Image processing was conducted using ImageJ software. The end of the cleavage product can be identified as either a 2’,3’-cyclic phosphate or a 3’ phosphate by observing the change in electrophoretic mobility of the product due to the loss of a negative charge by CIP.

#### RNA binding assay

1 μM dsRNA44 was incubated with 5 μM H234A in an RNA binding buffer [50 mM Tris-HCl (pH 8.0 at 25°C), 100 mM NaCl, and zero or 0.5 mM MnCl_2_ or CoCl_2_] at 37°C for 10 min. Subsequently, the sample was loaded onto a Superdex 200 Increase 10/300 GL column pre-equilibrated with the RNA binding buffer for gel filtration chromatography. The collected samples were further analyzed by 12.5 % SDS-PAGE with Coomassie blue staining and 20% TBE PAGE with ethidium bromide staining. RNA was visualized with a UVsolo Touch system (Analytik Jena, Germany). Image processing was conducted using ImageJ software.

#### Determination of the impact of the metal ions on nsp15 protein Differential scanning fluorimetry (DSF)

25 μL of sample containing 50 mM Tris-HCl (pH 7.4 at 25°C), 140 mM KCl, 0.5 mM EDTA, MnCl_2_, CoCl_2_, NiCl_2_, ZnCl_2_, or CuCl_2_, 5× SYPRO Orange dye (Sigma-Aldrich, USA) and 2 μM nsp15 was loaded into a 96-well PCR plate. DSF experiments were performed using a CFX Connect qPCR system. After an initial equilibration step at 25°C for 10 min, the temperature was increased by 0.5°C every 30 s until it reached 95°C. Fluorescence intensity was measured after each cycle using the FRET channel. The final data were analyzed using Bio-Rad CFX Maestro software. Each sample produced a melt curve, and the first derivative of the melt curve produced a peak, which provides the melting temperature (*T*_m_).

#### Native-PAGE

10 μL of sample containing 50 mM Tris-HCl (pH 7.4 at 25°C), 140 mM KCl, zero or 0.5 mM MnCl_2_, CoCl_2_, NiCl_2_, ZnCl_2_, or CuCl_2_, and 6 μM nsp15 was incubated at 37°C for 10 min. Then, 1 μL of 50 mM EDTA was added into the sample, followed by 10 min incubation at 37°C. After that, 3.7 μL of 4× Native-PAGE loading buffer (Solarbio, China) was added into the mixture. The samples were analyzed by 7.5% Native-PAGE with Coomassie blue staining. Protein size was determined with High Molecular Weight Native Electrophoresis Protein Marker II (Real-Times, China) in Native-PAGE analysis. In addition, the sample with CuCl_2_ was also analyzed by 12.5% SDS-PAGE.

#### Centrifugal analysis

20 μL of sample containing 50 mM Tris-HCl (pH 8.0 at 25°C), 100 mM NaCl, zero or 0.5 mM MnCl_2_, CoCl_2_, NiCl_2_, or ZnCl_2_, and 12 μM nsp15 was incubated at 37°C for 10 min. Then, 5 μL of the sample was extracted for SDS-PAGE. The remaining 15 μL of the sample was subjected to centrifugation at 13,000 rpm for 10 min. Subsequently, 5 μL of the supernatant was extracted for SDS-PAGE.

#### Negative-staining electron microscopy

A 3 µL sample of 0.03 mg/mL H234A, supplemented with 5 mM CoCl_2_, NiCl_2_, or ZnCl_2_, was applied to a copper grid (Beijing XXBR Technology, no. T10023), incubated for 1 min and absorbed with filter paper. The grid was then stained with 3 µL of 2% Uranium acetate for 1 min and air-dried. Images were acquired using a Talos L120C G2 120 kV transmission electron microscope.

#### Cryo-EM imaging and data processing

In our structural studies, we utilized the catalytically impaired H234A mutant, which was diluted to a final concentration of 7.5 µM. The nsp15/dsRNA complex sample was prepared by mixing H234A with dsRNA44 at a molar ratio of 1:10, followed by incubation on ice for 30 min in the presence of 5 mM Co^2+^. A 3 μL sample was applied to glow-discharged Quantifoil R1.2/1.3 300-mesh copper grid using a Vitrobot Mark IV (FEI) under the following conditions: blotting force of 3, blotting time of 3 s, 100% humidity, and a temperature of 8 °C. The grid was then plunge-frozen in liquid ethane and transferred to liquid nitrogen for storage. Initial screening was conducted on an FEI Glacios microscope (Core Facility of Wuhan University) operating at 200 kV and equipped with a Ceta D CMOS camera. A grid exhibiting optimal ice thickness and particle distribution was selected for high-resolution cryo-EM data collection. Data acquisition was performed using an FEI Titan Krios G4 microscope (Core Facility of Wuhan University) operating at 300 kV and equipped with a Gatan K3 direct electron detector. For the nsp15/dsRNA complex, a total of 4,868 movies were collected using Thermo Scientific EPU software in counting mode. Each movie consisted of 40 frames, with a cumulative electron dose of 50 electrons per Å2 and a pixel size of 0.84 Å. All datasets were processed using CryoSPARC.^65^

For the nsp15/dsRNA complex, the initial dataset underwent blob-picking followed by two rounds of 2D classification, yielding 319,161 particles for further processing. These particles were subjected to ab-initio reconstruction, followed by iterative rounds of homogeneous and heterogeneous refinement under C1 symmetry. The optimal class from this process was selected for additional homogeneous refinement with C1 symmetry, resulting in 244,604 particles. Subsequent global CTF refinement and additional homogeneous refinement were performed with C1 symmetry, followed by local CTF refinement and volume flipping. The flipped volume was then used for both homogeneous and non-uniform refinement under C1 symmetry, yielding a 6:1 nsp15/dsRNA map at 2.6 Å resolution. This 6:1 nsp15/dsRNA map was subsequently used for heterogeneous refinement under C1 symmetry. The best class obtained from this step underwent further homogeneous refinement under C1 symmetry. The refined volume was then combined with the initial 319,161 particles for another round of heterogeneous refinement. The optimal class from this final refinement step was subjected to homogeneous and non-uniform refinement under C1 symmetry, ultimately producing a 6:2 nsp15/dsRNA map at 2.78 Å resolution.

#### Model building and structure refinement

The 7TJ2 structure served as the initial model for constructing the 6:1 nsp15/dsRNA complex, which was subsequently used to build the 6:2 nsp15/dsRNA complex. All models were generated using Coot.^66^ Model refinement was performed through real-space refinement in PHENIX.^67^ The overall model quality was evaluated using MolProbity.^68^ Detailed statistical analyses are provided in Table S3. Structural figures were prepared using PyMOL and UCSF ChimeraX.^69^

#### Structural prediction of metal-bound nsp15 complexes by AlphaFold3

The structures of Mn^2+^-bound and Co^2+^-bound nsp15 complexes were predicted using AlphaFold3.^70^ The amino acid sequence of SARS-CoV-2 nsp15 was submitted to the AlphaFold Server (https://alphafoldserver.com/) with copies being six. Mn^2+^ and Co^2+^ were selected in two independent jobs, respectively, with copies being six. The predicted structures were analyzed using PyMOL, and the model ranks first among the output models (model 0) for each structure is showed.

## ACKNOWLEDGEMENTS

We thank all lab members for helpful discussion and technical assistance. We are grateful to Danyang Li and Xiangning Li at the Core Facility of Wuhan University for Cryo-EM data collection services. This work was supported by the National Natural Science Foundation of China (grant 32150009), the Feng Foundation of Biomedical Research, the Billionhome Venture Capital to B.Z., and National Key R&D Program of China (grant 2022YFA0912200) to L.W..

## AUTHOR CONTRIBUTIONS

X.L.W. and B.Z. conceived the project. B.Z. and L.F.W. acquired the funding and supervised the work. X.L.W. carried out all biochemical and molecular experiments unless otherwise indicated. J.L. performed negative-staining electron microscopy, cryo-EM imaging, data processing, model building, and structure refinement. Z.C.L. assisted with the protein expression and purification. X.L.W., J.L., L.F.W., and B.Z. analyzed the data. X.L.W., J.L., L.F.W., and B.Z. wrote the manuscript.

## SUPPLEMENTAL INFORMATION

Supplemental information can be found online.

## REFERENCES

1. V’Kovski, P., Kratzel, A., Steiner, S., Stalder, H., and Thiel, V. (2021). Coronavirus biology and replication: implications for SARS-CoV-2. Nat Rev Microbiol 19, 155–170. 10.1038/s41579-020-00468-6.

2. Proudfoot, A.E. (2002). Chemokine receptors: multifaceted therapeutic targets. Nature reviews. Immunology 2, 106–115. 10.1038/nri722.

3. Lazear, H.M., Schoggins, J.W., and Diamond, M.S. (2019). Shared and Distinct Functions of Type I and Type III Interferons. Immunity 50, 907–923. 10.1016/j.immuni.2019.03.025.

4. Blanco-Melo, D., Nilsson-Payant, B.E., Liu, W.-C., Uhl, S., Hoagland, D., Møller, R., Jordan, T.X., Oishi, K., Panis, M., Sachs, D., et al. (2020). Imbalanced Host Response to SARS-CoV-2 Drives Development of COVID-19. Cell 181, 1036–1045.e1039. 10.1016/j.cell.2020.04.026.

5. Channappanavar, R., Fehr, A.R., Vijay, R., Mack, M., Zhao, J., Meyerholz, D.K., and Perlman, S. (2016). Dysregulated Type I Interferon and Inflammatory Monocyte-Macrophage Responses Cause Lethal Pneumonia in SARS-CoV-Infected Mice. Cell host & microbe 19, 181–193. 10.1016/j.chom.2016.01.007.

6. Deng, X., and Baker, S.C. (2018). An "Old" protein with a new story: Coronavirus endoribonuclease is important for evading host antiviral defenses. Virology 517, 157–163. 10.1016/j.virol.2017.12.024.

7. Schneider, W.M., Chevillotte, M.D., and Rice, C.M. (2014). Interferon-stimulated genes: a complex web of host defenses. Annu Rev Immunol 32, 513–545. 10.1146/annurev-immunol-032713-120231.

8. Snijder, E.J., Bredenbeek, P.J., Dobbe, J.C., Thiel, V., Ziebuhr, J., Poon, L.L., Guan, Y., Rozanov, M., Spaan, W.J., and Gorbalenya, A.E. (2003). Unique and conserved features of genome and proteome of SARS-coronavirus, an early split-off from the coronavirus group 2 lineage. J Mol Biol 331, 991–1004. 10.1016/s0022-2836(03)00865-9.

9. Bhardwaj, K., Sun, J., Holzenburg, A., Guarino, L.A., and Kao, C.C. (2006). RNA recognition and cleavage by the SARS coronavirus endoribonuclease. J Mol Biol 361, 243–256. 10.1016/j.jmb.2006.06.021.

10. Deng, X., Hackbart, M., Mettelman, R.C., O’Brien, A., Mielech, A.M., Yi, G., Kao, C.C., and Baker, S.C. (2017). Coronavirus nonstructural protein 15 mediates evasion of dsRNA sensors and limits apoptosis in macrophages. Proc Natl Acad Sci U S A 114, E4251–E4260. 10.1073/pnas.1618310114.

11. Kindler, E., Gil-Cruz, C., Spanier, J., Li, Y., Wilhelm, J., Rabouw, H.H., Zust, R., Hwang, M., V’Kovski, P., Stalder, H., et al. (2017). Early endonuclease-mediated evasion of RNA sensing ensures efficient coronavirus replication. PLoS Pathog 13, e1006195. 10.1371/journal.ppat.1006195.

12. Comar, C.E., Otter, C.J., Pfannenstiel, J., Doerger, E., Renner, D.M., Tan, L.H., Perlman, S., Cohen, N.A., Fehr, A.R., and Weiss, S.R. (2022). MERS-CoV endoribonuclease and accessory proteins jointly evade host innate immunity during infection of lung and nasal epithelial cells. Proc Natl Acad Sci U S A 119, e2123208119. 10.1073/pnas.2123208119.

13. Otter, C.J., Bracci, N., Parenti, N.A., Ye, C., Asthana, A., Blomqvist, E.K., Tan, L.H., Pfannenstiel, J.J., Jackson, N., Fehr, A.R., et al. (2024). SARS-CoV-2 nsp15 endoribonuclease antagonizes dsRNA-induced antiviral signaling. Proc Natl Acad Sci U S A 121, e2320194121. 10.1073/pnas.2320194121.

14. Wu, Y., Zhang, H., Shi, Z., Chen, J., Li, M., Shi, H., Shi, D., Guo, L., and Feng, L. (2020). Porcine Epidemic Diarrhea Virus nsp15 Antagonizes Interferon Signaling by RNA Degradation of TBK1 and IRF3. Viruses 12, 599. 10.3390/v12060599.

15. Zhang, D., Ji, L., Chen, X., He, Y., Sun, Y., Ji, L., Zhang, T., Shen, Q., Wang, X., Wang, Y., et al. (2023). SARS-CoV-2 Nsp15 suppresses type I interferon production by inhibiting IRF3 phosphorylation and nuclear translocation. iScience 26. 10.1016/j.isci.2023.107705.

16. Chiu, H.P., Yeo, Y.Y., Lai, T.Y., Hung, C.T., Kowdle, S., Haas, G.D., Jiang, S., Sun, W., and Lee, B. (2024). SARS-CoV-2 Nsp15 antagonizes the cGAS-STING-mediated antiviral innate immune responses. bioRxiv : the preprint server for biology. 10.1101/2024.09.05.611469.

17. Wang, X., and Zhu, B. (2024). SARS-CoV-2 nsp15 preferentially degrades AU-rich dsRNA via its dsRNA nickase activity. Nucleic Acids Research 52, 5257–5272. 10.1093/nar/gkae290.

18. Ancar, R., Li, Y., Kindler, E., Cooper, D.A., Ransom, M., Thiel, V., Weiss, S.R., Hesselberth, J.R., and Barton, D.J. (2020). Physiologic RNA targets and refined sequence specificity of coronavirus EndoU. RNA 26, 1976–1999. 10.1261/rna.076604.120.

19. Hackbart, M., Deng, X., and Baker, S.C. (2020). Coronavirus endoribonuclease targets viral polyuridine sequences to evade activating host sensors. Proc Natl Acad Sci U S A 117, 8094–8103. 10.1073/pnas.1921485117.

20. Heusipp, G., Grotzinger, C., Herold, J., Siddell, S.G., and Ziebuhr, J. (1997). Identification and subcellular localization of a 41 kDa, polyprotein 1ab processing product in human coronavirus 229E-infected cells. J Gen Virol 78 ( *Pt 11*), 2789–2794. 10.1099/0022-1317-78-11-2789.

21. Shi, S.T., Schiller, J.J., Kanjanahaluethai, A., Baker, S.C., Oh, J.W., and Lai, M.M. (1999). Colocalization and membrane association of murine hepatitis virus gene 1 products and De novo-synthesized viral RNA in infected cells. J Virol 73, 5957–5969. 10.1128/JVI.73.7.5957-5969.1999.

22. Athmer, J., Fehr, A.R., Grunewald, M., Smith, E.C., Denison, M.R., and Perlman, S. (2017). In Situ Tagged nsp15 Reveals Interactions with Coronavirus Replication/Transcription Complex-Associated Proteins. mBio 8. 10.1128/mBio.02320-16.

23. Bhardwaj, K., Guarino, L., and Kao, C.C. (2004). The severe acute respiratory syndrome coronavirus Nsp15 protein is an endoribonuclease that prefers manganese as a cofactor. J Virol 78, 12218–12224. 10.1128/JVI.78.22.12218-12224.2004.

24. Ivanov, K.A., Hertzig, T., Rozanov, M., Bayer, S., Thiel, V., Gorbalenya, A.E., and Ziebuhr, J. (2004). Major genetic marker of nidoviruses encodes a replicative endoribonuclease. Proc Natl Acad Sci U S A 101, 12694–12699. 10.1073/pnas.0403127101.

25. Zhang, L., Li, L., Yan, L., Ming, Z., Jia, Z., Lou, Z., and Rao, Z. (2018). Structural and Biochemical Characterization of Endoribonuclease Nsp15 Encoded by Middle East Respiratory Syndrome Coronavirus. J Virol 92, e00893–00818. 10.1128/JVI.00893-18.

26. Kim, Y., Wower, J., Maltseva, N., Chang, C., Jedrzejczak, R., Wilamowski, M., Kang, S., Nicolaescu, V., Randall, G., Michalska, K., and Joachimiak, A. (2021). Tipiracil binds to uridine site and inhibits Nsp15 endoribonuclease NendoU from SARS-CoV-2. Commun Biol 4, 193. 10.1038/s42003-021-01735-9.

27. Godoy, A.S., Nakamura, A.M., Douangamath, A., Song, Y., Noske, G.D., Gawriljuk, V.O., Fernandes, R.S., Pereira, Humberto D M., Oliveira, Ketllyn Irene Z., Fearon, D., et al. (2023). Allosteric regulation and crystallographic fragment screening of SARS-CoV-2 NSP15 endoribonuclease. Nucleic Acids Research 51, 5255–5270. 10.1093/nar/gkad314.

28. Huang, T., Snell, K.C., Kalia, N., Gardezi, S., Guo, L., and Harris, M.E. (2023). Kinetic analysis of RNA cleavage by coronavirus Nsp15 endonuclease: Evidence for acid-base catalysis and substrate-dependent metal ion activation. J Biol Chem 299, 104787. 10.1016/j.jbc.2023.104787.

29. Bowman, A.B., and Aschner, M. (2014). Considerations on manganese (Mn) treatments for in vitro studies. Neurotoxicology 41, 141–142. 10.1016/j.neuro.2014.01.010.

30. Chen, P., Bornhorst, J., and Aschner, M. (2018). Manganese metabolism in humans. Front Biosci (Landmark Ed) 23, 1655–1679. 10.2741/4665.

31. Guarino, L.A., Bhardwaj, K., Dong, W., Sun, J., Holzenburg, A., and Kao, C. (2005). Mutational analysis of the SARS virus Nsp15 endoribonuclease: identification of residues affecting hexamer formation. J Mol Biol 353, 1106–1117. 10.1016/j.jmb.2005.09.007.

32. Xu, X., Zhai, Y., Sun, F., Lou, Z., Su, D., Xu, Y., Zhang, R., Joachimiak, A., Zhang, X.C., Bartlam, M., and Rao, Z. (2006). New Antiviral Target Revealed by the Hexameric Structure of Mouse Hepatitis Virus Nonstructural Protein nsp15. Journal of Virology 80, 7909–7917. 10.1128/jvi.00525-06.

33. Kim, Y., Jedrzejczak, R., Maltseva, N.I., Wilamowski, M., Endres, M., Godzik, A., Michalska, K., and Joachimiak, A. (2020). Crystal structure of Nsp15 endoribonuclease NendoU from SARS-CoV-2. Protein Sci 29, 1596–1605. 10.1002/pro.3873.

34. Frazier, M.N., Dillard, L.B., Krahn, J.M., Perera, L., Williams, J.G., Wilson, I.M., Stewart, Z.D., Pillon, M.C., Deterding, L.J., Borgnia, M.J., and Stanley, R.E. (2021). Characterization of SARS2 Nsp15 nuclease activity reveals it’s mad about U. Nucleic Acids Res 49, 10136–10149. 10.1093/nar/gkab719.

35. Pillon, M.C., Frazier, M.N., Dillard, L.B., Williams, J.G., Kocaman, S., Krahn, J.M., Perera, L., Hayne, C.K., Gordon, J., Stewart, Z.D., et al. (2021). Cryo-EM structures of the SARS-CoV-2 endoribonuclease Nsp15 reveal insight into nuclease specificity and dynamics. Nat Commun 12, 636. 10.1038/s41467-020-20608-z.

36. 36. Nedialkova, D.D., Ulferts, R., van den Born, E., Lauber, C., Gorbalenya, A.E., Ziebuhr, J., and Snijder, E.J. (2009). Biochemical Characterization of Arterivirus Nonstructural Protein 11 Reveals the Nidovirus-Wide Conservation of a Replicative Endoribonuclease. Journal of Virology 83, 5671–5682. 10.1128/jvi.00261-09.

37. Raines, R.T. (1998). Ribonuclease A. Chemical reviews 98, 1045–1066. 10.1021/cr960427h.

38. Hearon, J.Z. (1948). The configuration of cobaltodihistidine and oxy-bis (cobaltodihistidine). Journal of the National Cancer Institute 9, 1–11.

39. Ricagno, S., Egloff, M.P., Ulferts, R., Coutard, B., Nurizzo, D., Campanacci, V., Cambillau, C., Ziebuhr, J., and Canard, B. (2006). Crystal structure and mechanistic determinants of SARS coronavirus nonstructural protein 15 define an endoribonuclease family. Proc Natl Acad Sci U S A 103, 11892–11897. 10.1073/pnas.0601708103.

40. Mushegian, A., Sorokina, I., Eroshkin, A., and Dlakić, M. (2020). An ancient evolutionary connection between Ribonuclease A and EndoU families. Rna 26, 803–813. 10.1261/rna.074385.119.

41. Frazier, M.N., Wilson, I.M., Krahn, J.M., Butay, K.J., Dillard, L.B., Borgnia, M.J., and Stanley, R.E. (2022). Flipped over U: structural basis for dsRNA cleavage by the SARS-CoV-2 endoribonuclease. Nucleic Acids Res 50, 8290–8301. 10.1093/nar/gkac589.

42. Ito, F., Yang, H., Zhou, Z.H., and Chen, X.S. (2024). Structural basis for polyuridine tract recognition by SARS-CoV-2 Nsp15. Protein & Cell 15, 547–552. 10.1093/procel/pwae009.

43. Kirchdoerfer, R.N., and Ward, A.B. (2019). Structure of the SARS-CoV nsp12 polymerase bound to nsp7 and nsp8 co-factors. Nat Commun 10, 2342. 10.1038/s41467-019-10280-3.

44. Yin, W., Mao, C., Luan, X., Shen, D.D., Shen, Q., Su, H., Wang, X., Zhou, F., Zhao, W., Gao, M., et al. (2020). Structural basis for inhibition of the RNA-dependent RNA polymerase from SARS-CoV-2 by remdesivir. Science (New York, N.Y.) 368, 1499–1504. 10.1126/science.abc1560.

45. Lin, S., Chen, H., Chen, Z., Yang, F., Ye, F., Zheng, Y., Yang, J., Lin, X., Sun, H., Wang, L., et al. (2021). Crystal structure of SARS-CoV-2 nsp10 bound to nsp14-ExoN domain reveals an exoribonuclease with both structural and functional integrity. Nucleic Acids Research 49, 5382–5392. 10.1093/nar/gkab320.

46. Newman, J.A., Douangamath, A., Yadzani, S., Yosaatmadja, Y., Aimon, A., Brandão-Neto, J., Dunnett, L., Gorrie-stone, T., Skyner, R., Fearon, D., et al. (2021). Structure, mechanism and crystallographic fragment screening of the SARS-CoV-2 NSP13 helicase. Nature Communications 12. 10.1038/s41467-021-25166-6.

47. Moeller, N.H., Shi, K., Demir, Ö., Belica, C., Banerjee, S., Yin, L., Durfee, C., Amaro, R.E., and Aihara, H. (2022). Structure and dynamics of SARS-CoV-2 proofreading exoribonuclease ExoN. Proceedings of the National Academy of Sciences 119. 10.1073/pnas.2106379119.

48. Domingo, J.L., and Marquès, M. (2021). The effects of some essential and toxic metals/metalloids in COVID-19: A review. Food and Chemical Toxicology 152. 10.1016/j.fct.2021.112161.

49. Ni, Y.-Q., Zeng, H.-H., Song, X.-W., Zheng, J., Wu, H.-Q., Liu, C.-T., and Zhang, Y. (2022). Potential metal-related strategies for prevention and treatment of COVID-19. Rare Metals 41, 1129–1141. 10.1007/s12598-021-01894-y.

50. Tabatabaeizadeh, S.-A. (2022). Zinc supplementation and COVID-19 mortality: a meta-analysis. European Journal of Medical Research 27. 10.1186/s40001-022-00694-z.

51. Khan, K.M., Zimpfer, M.J., Sultana, R., Parvez, T.M., Navas-Acien, A., and Parvez, F. (2023). Role of Metals on SARS-CoV-2 Infection: a Review of Recent Epidemiological Studies. Current Environmental Health Reports 10, 353–368. 10.1007/s40572-023-00409-4.

52. Maret, W. (2015). Analyzing free zinc(II) ion concentrations in cell biology with fluorescent chelating molecules. Metallomics : integrated biometal science 7, 202–211. 10.1039/c4mt00230j.

53. Outten, C.E., and O’Halloran, T.V. (2001). Femtomolar sensitivity of metalloregulatory proteins controlling zinc homeostasis. Science (New York, N.Y.) 292, 2488–2492. 10.1126/science.1060331.

54. Vinkenborg, J.L., Nicolson, T.J., Bellomo, E.A., Koay, M.S., Rutter, G.A., and Merkx, M. (2009). Genetically encoded FRET sensors to monitor intracellular Zn2+ homeostasis. Nature methods 6, 737–740. 10.1038/nmeth.1368.

55. Qin, Y., Dittmer, P.J., Park, J.G., Jansen, K.B., and Palmer, A.E. (2011). Measuring steady-state and dynamic endoplasmic reticulum and Golgi Zn2+ with genetically encoded sensors. Proc Natl Acad Sci U S A 108, 7351–7356. 10.1073/pnas.1015686108.

56. Kambe, T., Tsuji, T., Hashimoto, A., and Itsumura, N. (2015). The Physiological, Biochemical, and Molecular Roles of Zinc Transporters in Zinc Homeostasis and Metabolism. Physiological reviews 95, 749–784. 10.1152/physrev.00035.2014.

57. Kimura, T., and Kambe, T. (2016). The Functions of Metallothionein and ZIP and ZnT Transporters: An Overview and Perspective. International Journal of Molecular Sciences 17. 10.3390/ijms17030336.

58. Perry, D.K., Smyth, M.J., Stennicke, H.R., Salvesen, G.S., Duriez, P., Poirier, G.G., and Hannun, Y.A. (1997). Zinc is a potent inhibitor of the apoptotic protease, caspase-3. A novel target for zinc in the inhibition of apoptosis. J Biol Chem 272, 18530–18533. 10.1074/jbc.272.30.18530.

59. Stennicke, H.R., and Salvesen, G.S. (1997). Biochemical characteristics of caspases-3, -6, -7, and -8. J Biol Chem 272, 25719–25723. 10.1074/jbc.272.41.25719.

60. Chai, F., Truong-Tran, A.Q., Evdokiou, A., Young, G.P., and Zalewski, P.D. (2000). Intracellular zinc depletion induces caspase activation and p21 Waf1/Cip1 cleavage in human epithelial cell lines. The Journal of infectious diseases 182 *Suppl 1*, S85–92. 10.1086/315914.

61. Summersgill, H., England, H., Lopez-Castejon, G., Lawrence, C.B., Luheshi, N.M., Pahle, J., Mendes, P., and Brough, D. (2014). Zinc depletion regulates the processing and secretion of IL-1β. Cell death & disease 5, e1040. 10.1038/cddis.2013.547.

62. Du, M., and Chen, Z.J. (2018). DNA-induced liquid phase condensation of cGAS activates innate immune signaling. Science (New York, N.Y.) 361, 704–709. 10.1126/science.aat1022.

63. Henry, T., Gong, X., Gu, W., Fu, S., Zou, G., and Jiang, Z. (2024). Zinc homeostasis regulates caspase activity and inflammasome activation. PLOS Pathogens 20. 10.1371/journal.ppat.1012805.

64. 64. Leal, K.N.d.S., Santos da Silva, A.B., Fonseca, E.K.B., Moreira, O.B.d.O., de Lemos, L.M., Leal de Oliveira, M.A., Stewart, A.J., and Arruda, M.A.Z. (2024). Metallomic analysis of urine from individuals with and without Covid-19 infection reveals extensive alterations in metal homeostasis. Journal of Trace Elements in Medicine and Biology 86. 10.1016/j.jtemb.2024.127557.

65. Punjani, A., Rubinstein, J.L., Fleet, D.J., and Brubaker, M.A. (2017). cryoSPARC: algorithms for rapid unsupervised cryo-EM structure determination. Nature methods 14, 290–296. 10.1038/nmeth.4169.

66. Emsley, P., and Cowtan, K. (2004). Coot: model-building tools for molecular graphics. Acta crystallographica. Section D, Biological crystallography 60, 2126–2132. 10.1107/s0907444904019158.

67. Liebschner, D., Afonine, P.V., Baker, M.L., Bunkóczi, G., Chen, V.B., Croll, T.I., Hintze, B., Hung, L.W., Jain, S., McCoy, A.J., et al. (2019). Macromolecular structure determination using X-rays, neutrons and electrons: recent developments in Phenix. Acta crystallographica. Section D, Structural biology 75, 861–877. 10.1107/s2059798319011471.

68. Williams, C.J., Headd, J.J., Moriarty, N.W., Prisant, M.G., Videau, L.L., Deis, L.N., Verma, V., Keedy, D.A., Hintze, B.J., Chen, V.B., et al. (2018). MolProbity: More and better reference data for improved all-atom structure validation. Protein Sci 27, 293–315. 10.1002/pro.3330.

69. Pettersen, E.F., Goddard, T.D., Huang, C.C., Meng, E.C., Couch, G.S., Croll, T.I., Morris, J.H., and Ferrin, T.E. (2021). UCSF ChimeraX: Structure visualization for researchers, educators, and developers. Protein Sci 30, 70–82. 10.1002/pro.3943.

70. Abramson, J., Adler, J., Dunger, J., Evans, R., Green, T., Pritzel, A., Ronneberger, O., Willmore, L., Ballard, A.J., Bambrick, J., et al. (2024). Accurate structure prediction of biomolecular interactions with AlphaFold 3. Nature 630, 493–500. 10.1038/s41586-024-07487-w.

