## Supplemental Data 1 for "Divalent cations govern the activity of coronavirus nsp15"

**Figure S1**

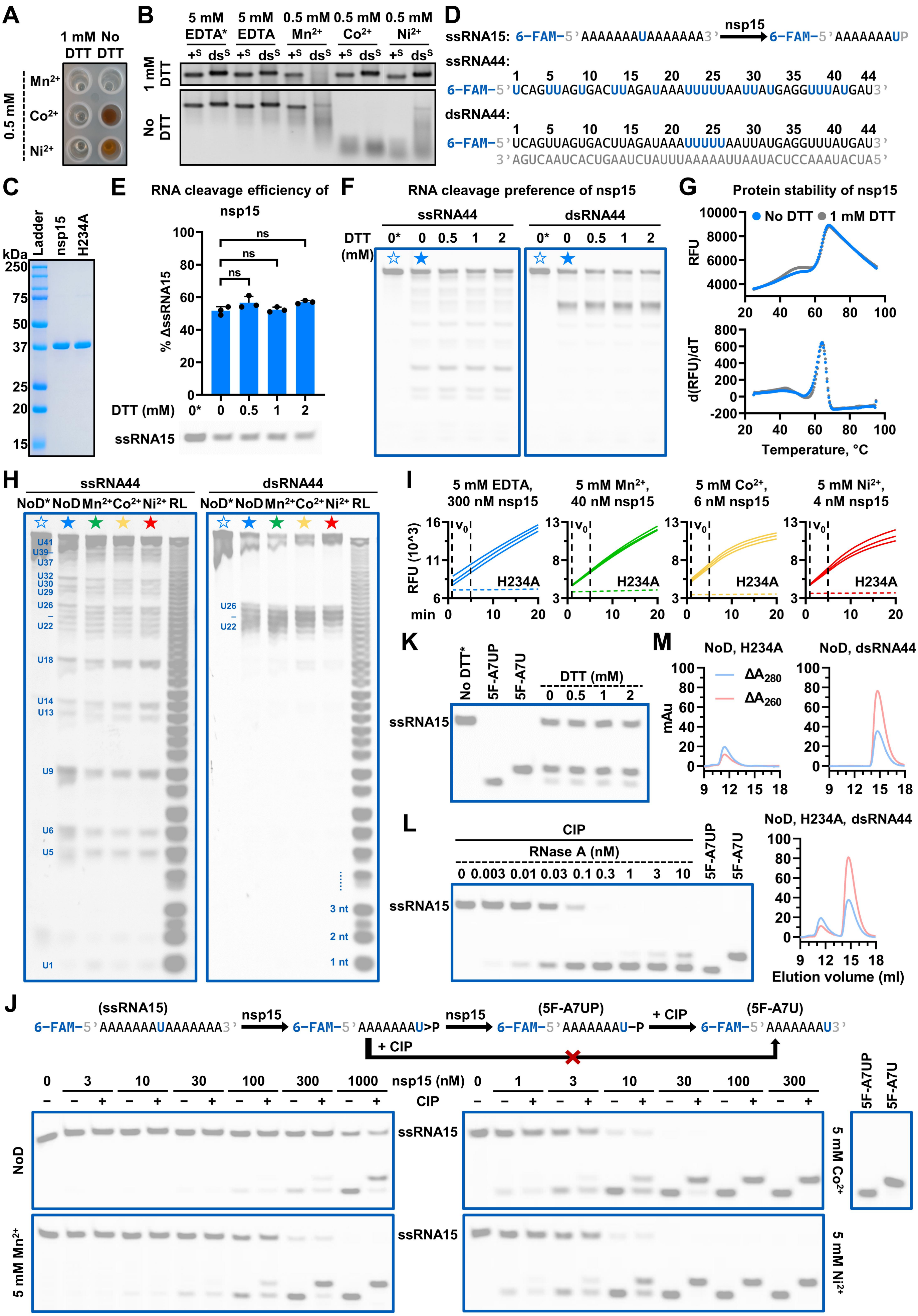

A
1 mM DTT
No DTT
0.5 mM
Mn2+
Co2+
Ni2+
B
5 mM EDTA*
5 mM EDTA
0.5 mM Mn2+
0.5 mM Co2+
0.5 mM Ni2+
+S dsS
1 mM DTT
No DTT
C
kDa
Ladder
nsp15
H234A
250
75
50
37
25
20
15
D
ssRNA15: 6-FAM-5'AAAAAAAUAAAAAAA3'
nsp15
6-FAM-5'AAAAAAAUP
ssRNA44:
1 5 10 15 20 25 30 35 40 44
6-FAM-5'UCAGUUAGUGACUUAGAUAAAUUUUUAAUUAUGAGGUUUAUGAU3'
dsRNA44:
1 5 10 15 20 25 30 35 40 44
6-FAM-5'UCAGUUAGUGACUUAGAUAAAUUUUUAAUUAUGAGGUUUAUGAU3'
3'AGUCAAUCACUGAAUCUAUUUAAAAAUUAAUACUCCAAAUACUA5'
E
RNA cleavage efficiency of nsp15
% ΔssRNA15
ns
DTT (mM) 0* 0 0.5 1 2
ssRNA15
F
RNA cleavage preference of nsp15
ssRNA44
dsRNA44
DTT (mM)
0* 0 0.5 1 2
G
Protein stability of nsp15
No DTT
1 mM DTT
RFU
d(RFU)/dT
Temperature, °C
H
ssRNA44
dsRNA44
NoD* NoD Mn2+ Co2+ Ni2+ RL
U41
U39
U37
U32
U30
U29
U26
U22
U18
U14
U13
U9
U6
U5
U1
3 nt
2 nt
1 nt
I
5 mM EDTA, 300 nM nsp15
5 mM Mn2+, 40 nM nsp15
5 mM Co2+, 6 nM nsp15
5 mM Ni2+, 4 nM nsp15
RFU (10^3)
v0
H234A
min
K
No DTT*
5F-A7UP
5F-A7U
DTT (mM)
0 0.5 1 2
ssRNA15
L
CIP
RNase A (nM)
0 0.003 0.01 0.03 0.1 0.3 1 3 10
5F-A7UP
5F-A7U
ssRNA15
M
NoD, H234A
NoD, dsRNA44
NoD, H234A, dsRNA44
ΔA280
ΔA260
mAu
Elution volume (ml)
J
(ssRNA15)
6-FAM-5'AAAAAAAUAAAAAAA3'
nsp15
6-FAM-5'AAAAAAAU>P
nsp15
(5F-A7UP)
6-FAM-5'AAAAAAAU-P
+ CIP
(5F-A7U)
6-FAM-5'AAAAAAAU3'
+ CIP
0 3 10 30 100 300 1000 nsp15 (nM)
0 1 3 10 30 100 300
CIP
NoD
5 mM Mn2+
ssRNA15
5 mM Co2+
5 mM Ni2+
5F-A7UP
5F-A7U

**Figure S1. $Co^{2+}$ and $Ni^{2+}$ exhibit an enhancing effect on nsp15 activity in the absence of DTT, related to Figure 1**

(A) $Co^{2+}$ and $Ni^{2+}$ were reduced by DTT, as evidenced by a color change from clear to red.

(B) Reduction of $Co^{2+}$ and $Ni^{2+}$ by DTT resulted in the elimination of their enhancing effect on the RNA cleavage activity of nsp15. $+^S$ and $ds^S$ refer to the positive-sense ssRNA form and the dsRNA form of a SARS-CoV-2 mini-genome, respectively.

(C) SDS-PAGE analysis of the purified wild-type SARS-CoV-2 nsp15 with an N-terminal 6× His tag (approximately 40 kDa) and its active-site mutant, H234A.

(D) Sequences of ssRNA15, ssRNA44, and dsRNA44. The preferred cleavage sites of nsp15 on these RNA substrates are indicated in blue.

(E) DTT exhibited no significant effect on the RNA cleavage efficiency of nsp15. In the absence of divalent metal ions, 1000 nM nsp15 was incubated with ssRNA15 under various concentrations of DTT. Reduction of the ssRNA15 substrate in every reaction was calculated as % ΔssRNA15. The average and standard deviation of three independent reactions are graphed. Student's t-test was performed. ns, not significant, $p > 0.05$.

(F) DTT exhibited no significant effect on the RNA cleavage preference of nsp15. In the absence of divalent metal ions, 40 nM nsp15 was incubated with ssRNA44 or dsRNA44 under various concentrations of DTT. The samples marked with pentagrams were used to analyze the preferred cleavage sites of nsp15 on ssRNA44 and dsRNA44 (see Figure S1H).

(G) DTT exhibited no significant effect on the protein stability of nsp15. In the DSF analysis, each sample produced a melt curve (as shown on the top), and the first derivative of the melt curve produced a peak (as shown on the bottom), which provides the melting temperature ($T_m$). RFU refers to relative fluorescence unit.

(H) Identification of the preferred cleavage sites of nsp15 on ssRNA44 and dsRNA44. NoD and RL refer to no divalent metal ions and RNA ladder, respectively.

(I) Cleavage of FRET-RNA by nsp15. The initial velocity ($v_0$) was estimated using the slope from linear regressions of the reaction monitored during 1–5 min.

(J) Detection of the ability of nsp15 to catalyze both transphosphorylation of RNA to form a 2',3'-cyclic phosphodiester and its subsequent hydrolysis to a 3'-phosphomonoester.

(K) DTT did not affect the hydrolysis of 2',3'-cyclic phosphodiesters by nsp15.

(L) Detection of the ability of RNase A to catalyze both transphosphorylation of RNA to form a 2',3'-cyclic phosphodiester and its subsequent hydrolysis to a 3'-phosphomonoester.

(M) Gel filtration chromatography results of H234A, dsRNA44, and the mixture of H234A and dsRNA44.

In (B, E, F, H, I, and K), the reactions containing the H234A mutant were used as negative controls, and these are marked with an asterisk (*).

#### Figure S2

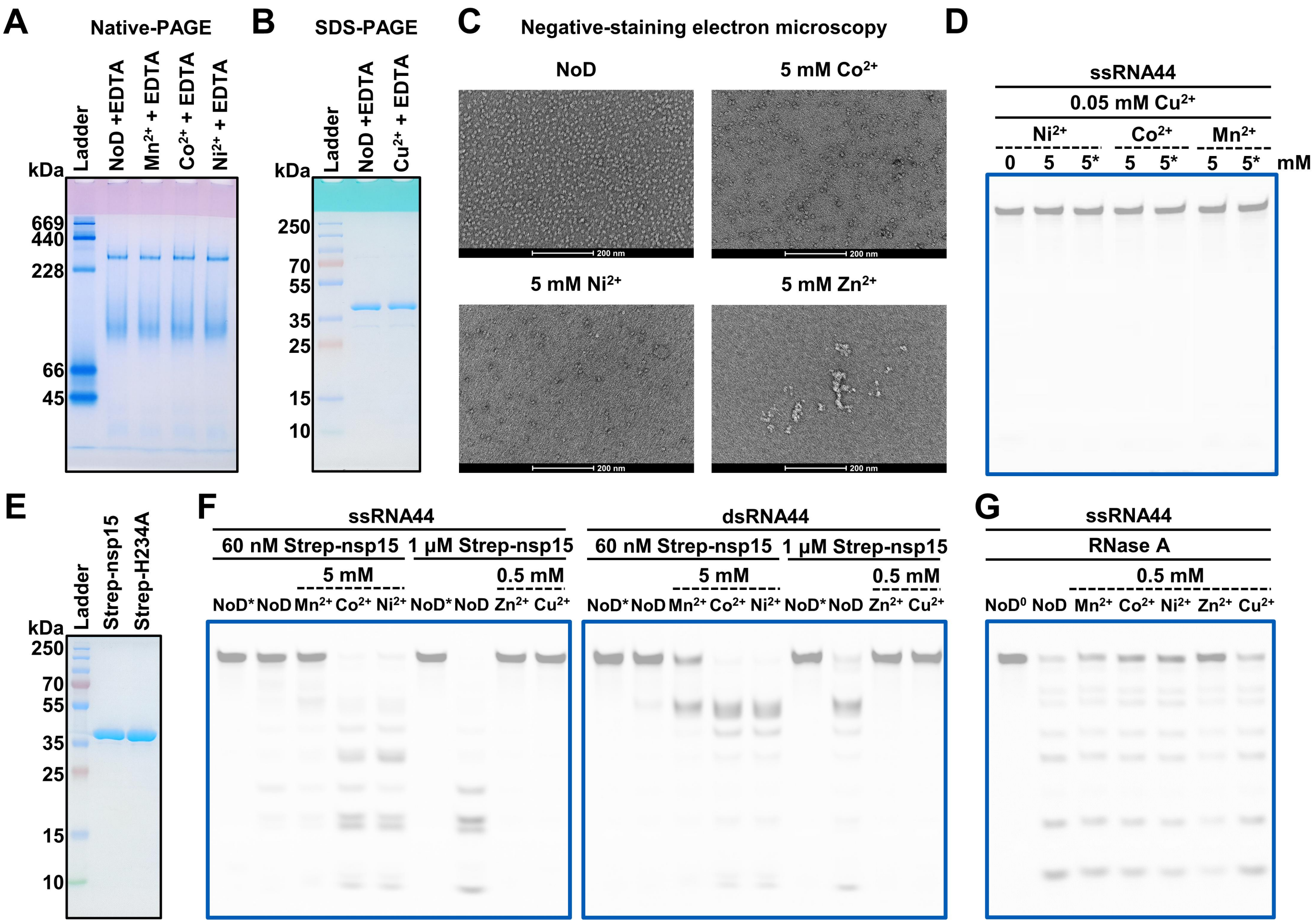

A
Native-PAGE
kDa
Ladder
NoD +EDTA
Mn2+ + EDTA
Co2+ + EDTA
Ni2+ + EDTA
669
440
228
66
45
B
SDS-PAGE
kDa
Ladder
NoD +EDTA
Cu2+ + EDTA
250
70
55
35
25
15
10
C
Negative-staining electron microscopy
NoD
5 mM Co2+
5 mM Ni2+
5 mM Zn2+
D
ssRNA44
0.05 mM Cu2+
Ni2+
Co2+
Mn2+
0
5
5*
5
5*
5
5*
mM
E
kDa
Ladder
Strep-nsp15
Strep-H234A
250
70
55
35
25
15
10
F
ssRNA44
dsRNA44
60 nM Strep-nsp15
1 μM Strep-nsp15
5 mM
0.5 mM
NoD*
NoD
Mn2+
Co2+
Ni2+
NoD*
NoD
Zn2+
Cu2+
G
ssRNA44
RNase A
0.5 mM
NoD0
NoD
Mn2+
Co2+
Ni2+
Zn2+
Cu2+

**Figure S2. Effects of $Mn^{2+}$, $Co^{2+}$, $Ni^{2+}$, $Zn^{2+}$ and $Cu^{2+}$ on nsp15 and RNase A, related to Figure 2**

(A) Native-PAGE analysis of the nsp15 protein pretreated with $Mn^{2+}$, $Co^{2+}$, and $Ni^{2+}$. The nsp15 hexamer has an estimated molecular weight of approximately 240 kDa.

(B) SDS-PAGE analysis of the nsp15 protein pretreated with $Cu^{2+}$.

(C) Representative negative-staining images of the H234A protein in the presence of $Co^{2+}$, $Ni^{2+}$, or $Zn^{2+}$.

(D) Cleavage of ssRNA44 by 20 nM nsp15 in the presence of $Ni^{2+}$, $Co^{2+}$, or $Mn^{2+}$, as well as in the simultaneous presence of $Cu^{2+}$.

(E) SDS-PAGE analysis of the purified wild-type SARS-CoV-2 nsp15 with an N-terminal Strep-Tag II (approximately 40 kDa) and its active-site mutant, H234A.

(F) Cleavage of ssRNA44 and dsRNA44 by Strep-nsp15 in the presence of $Mn^{2+}$, $Co^{2+}$, $Ni^{2+}$, $Zn^{2+}$ or $Cu^{2+}$.

(G) Cleavage of ssRNA44 by RNase A in the presence of $Mn^{2+}$, $Co^{2+}$, $Ni^{2+}$, $Zn^{2+}$ or $Cu^{2+}$. The samples lacking RNase A were used as negative controls, and these are marked with $^{0}$.

**Figure S3**

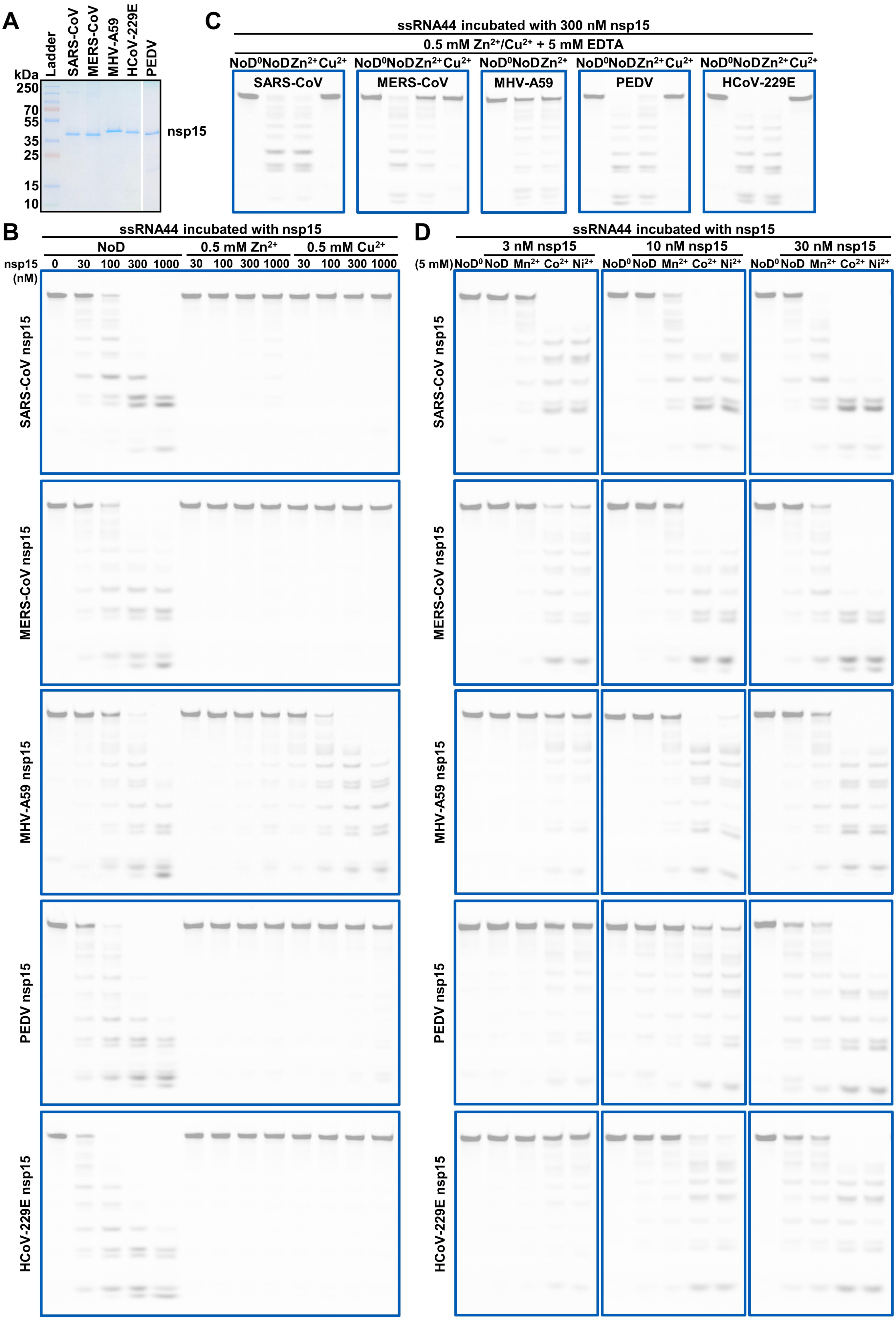
A
Ladder
SARS-CoV
MERS-CoV
MHV-A59
HCoV-229E
PEDV
kDa
250
70
55
35
25
15
10
nsp15
C
ssRNA44 incubated with 300 nM nsp15
0.5 mM Zn2+/Cu2+ + 5 mM EDTA
NoD0 NoD Zn2+ Cu2+
SARS-CoV
MERS-CoV
MHV-A59
PEDV
HCoV-229E
B
ssRNA44 incubated with nsp15
NoD
0.5 mM Zn2+
0.5 mM Cu2+
nsp15 (nM)
0 30 100 300 1000 30 100 300 1000 30 100 300 1000
SARS-CoV nsp15
MERS-CoV nsp15
MHV-A59 nsp15
PEDV nsp15
HCoV-229E nsp15
D
ssRNA44 incubated with nsp15
3 nM nsp15
10 nM nsp15
30 nM nsp15
(5 mM) NoD0 NoD Mn2+ Co2+ Ni2+
SARS-CoV nsp15
MERS-CoV nsp15
MHV-A59 nsp15
PEDV nsp15
HCoV-229E nsp15

**Figure S3. Specific effects of $Co^{2+}$, $Ni^{2+}$ and $Zn^{2+}$ on nsp15 are widespread in coronaviruses, related to Figure 3**

(A) SDS-PAGE analysis of the recombinant nsp15 proteins of various coronaviruses.

(B) Cleavage of ssRNA44 by nsp15 of various coronaviruses in the presence of $Zn^{2+}$ or $Cu^{2+}$.

(C) Cleavage of ssRNA44 by nsp15 of various coronaviruses, with pretreatment of the protein with $Zn^{2+}$ or $Cu^{2+}$.

(D) Cleavage of ssRNA44 by nsp15 of various coronaviruses in the presence of $Mn^{2+}$, $Co^{2+}$, or $Ni^{2+}$.

#### Figure S4

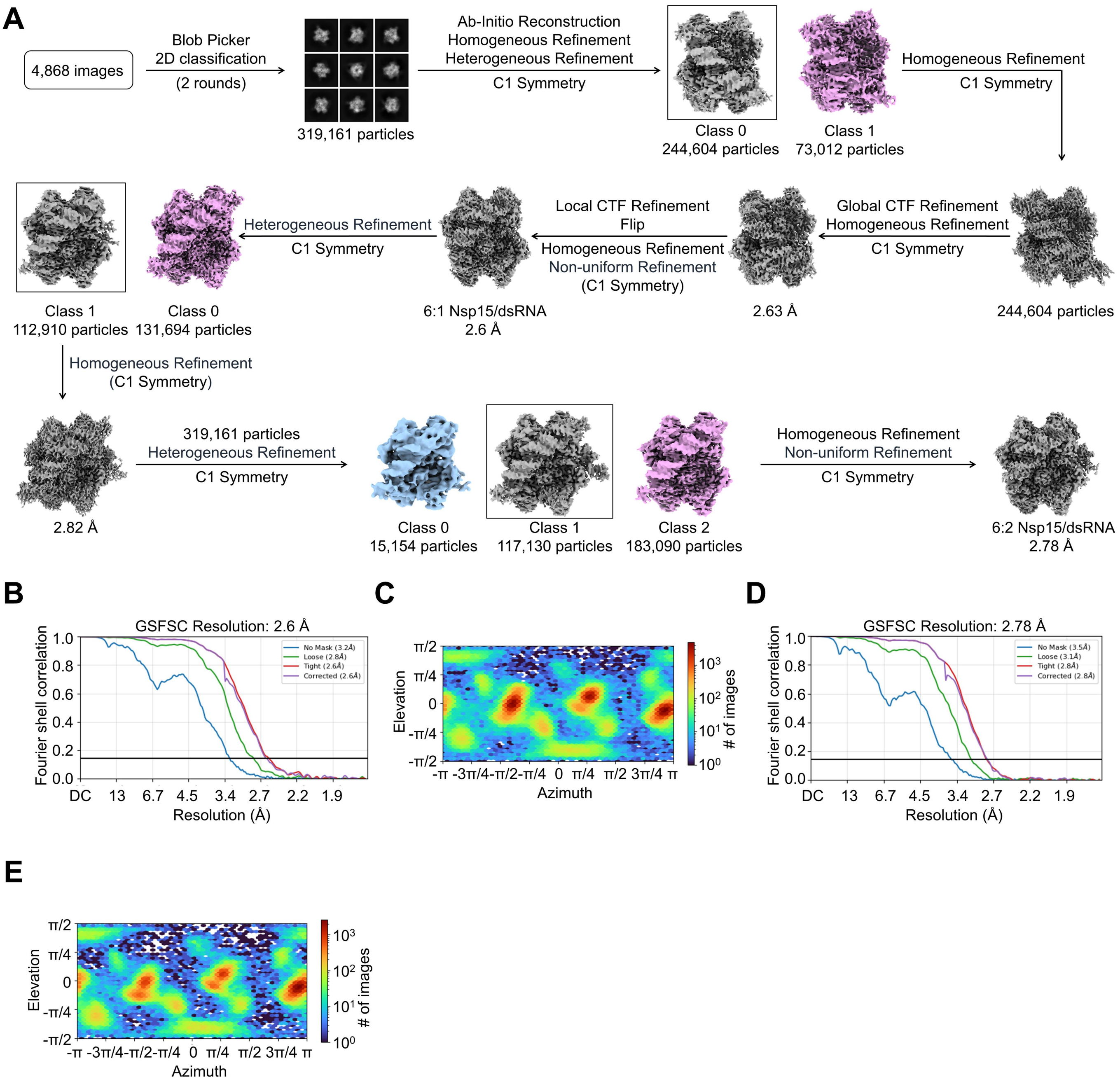
A
4,868 images
Blob Picker
2D classification
(2 rounds)
319,161 particles
Ab-Initio Reconstruction
Homogeneous Refinement
Heterogeneous Refinement
C1 Symmetry
Class 0
244,604 particles
Class 1
73,012 particles
Homogeneous Refinement
C1 Symmetry
244,604 particles
Global CTF Refinement
Homogeneous Refinement
C1 Symmetry
2.63 Å
Local CTF Refinement
Flip
Homogeneous Refinement
Non-uniform Refinement
(C1 Symmetry)
6:1 Nsp15/dsRNA
2.6 Å
Heterogeneous Refinement
C1 Symmetry
Class 1
112,910 particles
Class 0
131,694 particles
Homogeneous Refinement
(C1 Symmetry)
2.82 Å
319,161 particles
Heterogeneous Refinement
C1 Symmetry
Class 0
15,154 particles
Class 1
117,130 particles
Class 2
183,090 particles
Homogeneous Refinement
Non-uniform Refinement
C1 Symmetry
6:2 Nsp15/dsRNA
2.78 Å
B
GSFSC Resolution: 2.6 Å
No Mask (3.2Å)
Loose (2.8Å)
Tight (2.6Å)
Corrected (2.6Å)
Fourier shell correlation
Resolution (Å)
C
Elevation
Azimuth
### of images
D
GSFSC Resolution: 2.78 Å
No Mask (3.5Å)
Loose (3.1Å)
Tight (2.8Å)
Corrected (2.8Å)
Fourier shell correlation
Resolution (Å)
E
Elevation
Azimuth
### of images

**Figure S4. Reconstruction of the 6:1 and 6:2 nsp15/dsRNA complexes, related to Figure 4**

(A) Data processing workflow for the reconstruction of the 6:1 and 6:2 nsp15/dsRNA complexes.

(B) Gold-standard FSC curves of the 6:1 nsp15/dsRNA complex indicate an overall resolution of 2.6 Å at FSC=0.143.

(C) The orientation distribution plot of the 3D reconstruction of the 6:1 nsp15/dsRNA complex.

(D) Gold-standard FSC curves of the 6:2 nsp15/dsRNA complex indicate an overall resolution of 2.78 Å at FSC=0.143.

(E) The orientation distribution plot of the 3D reconstruction of the 6:2 nsp15/dsRNA complex.

Figure S5

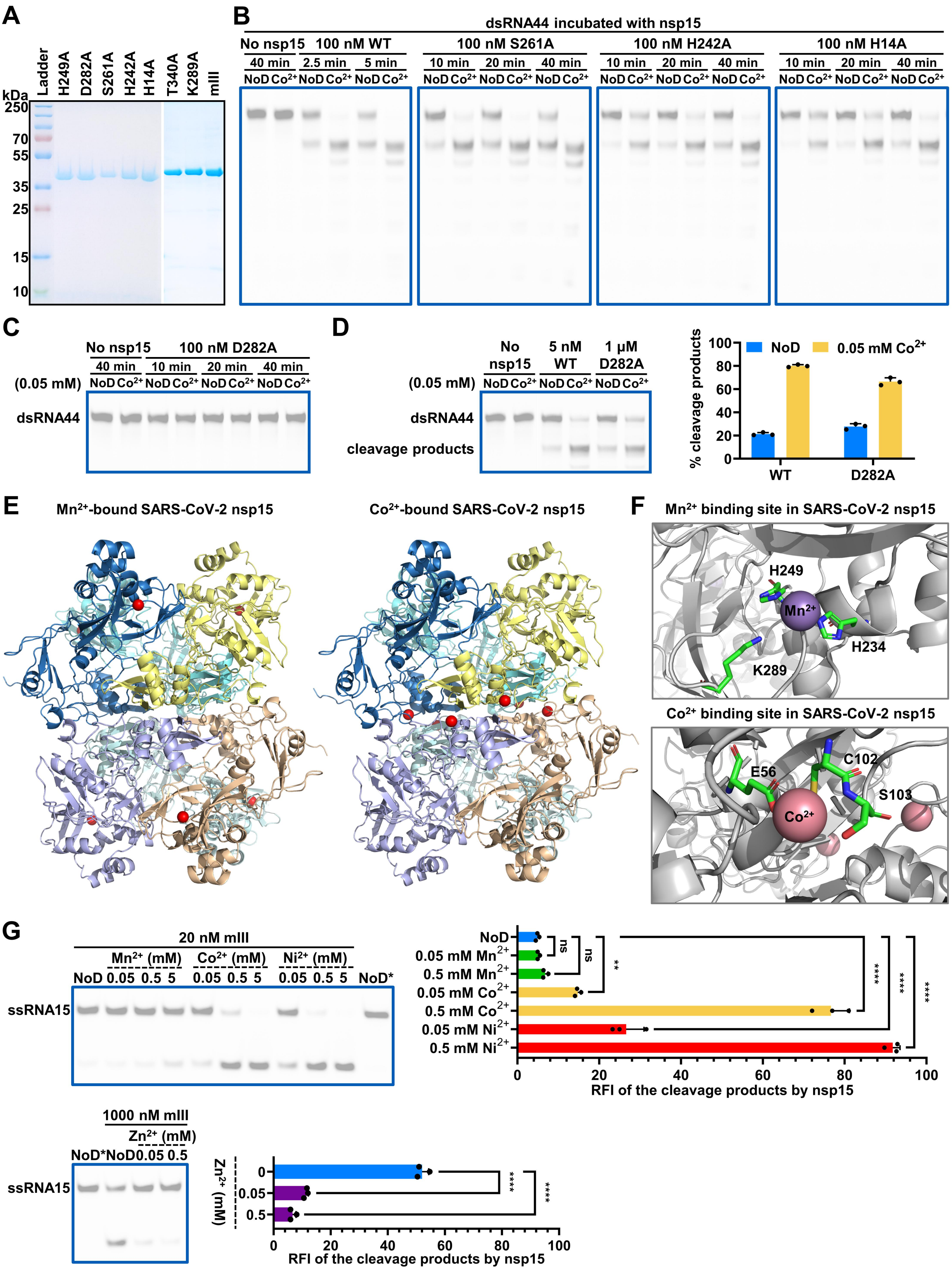
A
kDa
Ladder
H249A
D282A
S261A
H242A
H14A
T340A
K289A
mIII
250
70
55
35
25
15
10
B
dsRNA44 incubated with nsp15
No nsp15
100 nM WT
100 nM S261A
100 nM H242A
100 nM H14A
40 min
2.5 min
5 min
10 min
20 min
40 min
NoD Co2+
C
No nsp15
100 nM D282A
(0.05 mM)
dsRNA44
D
No nsp15
5 nM WT
1 μM D282A
cleavage products
% cleavage products
NoD
0.05 mM Co2+
WT
D282A
E
Mn2+-bound SARS-CoV-2 nsp15
Co2+-bound SARS-CoV-2 nsp15
F
Mn2+ binding site in SARS-CoV-2 nsp15
H249
Mn2+
H234
K289
Co2+ binding site in SARS-CoV-2 nsp15
E56
C102
S103
Co2+
G
20 nM mIII
Mn2+ (mM)
Co2+ (mM)
Ni2+ (mM)
NoD 0.05 0.5 5 0.05 0.5 5 0.05 0.5 5 NoD*
ssRNA15
NoD
0.05 mM Mn2+
0.5 mM Mn2+
0.05 mM Co2+
0.5 mM Co2+
0.05 mM Ni2+
0.5 mM Ni2+
ns
**
****
RFI of the cleavage products by nsp15
1000 nM mIII
Zn2+ (mM)
NoD* NoD 0.05 0.5
Zn2+ (mM)
RFI of the cleavage products by nsp15

**Figure S5. Impact of mutations in residues, excluding those within the active site, on the functions of divalent metal ions, related to Figure 5**

**(A)** SDS-PAGE analysis of the purified nsp15 mutants.

**(B, C, D)** Impact of mutations at S261, H242, H14, and D282, on the role of $Co^{2+}$. In **(D)**, the ratio of the gray value of the cleavage product band to the sum of the gray values of the remaining substrate band and the cleavage product band is defined as % cleavage products. The average and standard deviation of three independent reactions are graphed.

**(E)** Predicted structures of metal-bound nsp15 complexes by AlphaFold3. Metal ions are represented by red balls in the diagram.

**(F)** Enlarged view of the metal binding sites of nsp15 predicted by AlphaFold3. The prediction was made that $Mn^{2+}$ would bind at the active site, while $Co^{2+}$ was localized in the proximity of E56, C102, and S103.

**(G)** Quantification of the effects of $Mn^{2+}$, $Co^{2+}$, $Ni^{2+}$, and $Zn^{2+}$ on the EndoU activity of the nsp15 mutant with three point mutations including E56A, C102A, and S103A. The ratio of the gray value of the cleavage product band to the sum of the gray values of the remaining substrate band and the cleavage product band is defined as the relative fluorescence intensity (RFI) of the cleavage products. The average and standard deviation of three independent reactions are graphed. Student's t-test was performed. ns, not significant, $p > 0.05$; $^{**}p < 0.01$; $^{****}p < 0.0001$.

**Table S2. Primers for cloning, related to STAR Methods**

| Primers | Primer sequences (5' to 3') |
|---|---|
| F-Strep-tag II | TGGAGCCACCCGCAGTTCGAAAAGAGCAGCGGCAGCCTGG |
| R-Strep-tag II | CTTTTCGAACTGCGGGTGGCTCCAGCTGCTGCCCATGGTATATCTCC |
| F-H14A | AAAGGTGCGTTCGATGGTCAGCAAGGCGAGG |
| R-H14A | ATCGAACGCACCTTTGTTAACCACGTTAAACGC |
| F-H234A | TTTGAAGCGATCGTTTACGGCGATTTCAGCCATAG |
| R-H234A | AACGATCGCTTCAAACGCATAACCCTCCAGTTTGT |
| F-H242A | TTCAGCGCGAGCCAGCTGGGTGGCC |
| R-H242A | CTGGCTCGCGCTGAAATCGCCGTAAACGATGTGTTC |
| F-H249A | GGCCTGGCGCTGCTGATTGGTCTGGC |
| R-H249A | CAGCAGCGCCAGGCCACCCAGCTGGC |
| F-S261A | AAAGAGGCGCCGTTTGAGCTGGAAGATTTCATCCC |
| R-S261A | AAACGGCGCCTCTTTGAAACGCTTCGCCAGACC |
| F-D282A | ATTACCGCGGCGCAGACCGGCAGC |
| R-D282A | CTGCGCCGCGGTAATAAAATAGTTCTTCACGGTGC |
| F-K289A | AGCAGCGCGTGCGTGTGCAGCGTTATCGAC |
| R-K289A | CACGCACGCGCTGCTGCCGGTCTGC |
| F-T340A | GTGGAAGCGTTCTATCCGAAACTGCAATAAGAATTCGAGC |
| R-T340A | ATAGAACGCTTCCACGTGGCCGTCCTTGC |
| F-E56A | GCGTTCGCGCTGTGGGCGAAGCG |
| R-E56A | CCACAGCGCGAACGCAACGTTCACCG |
| F-C102A&S103A | CGTTGCGGCGATGACCGATATCGCGAAGAAACCG |
| R-C102A&S103A | GTCATCGCCGCAACGCCAATGGTGCTGATGTGC |

**Table S3. Cryo-EM data collection, refinement and validation statistics, related to Figure S4**

| | 6:1 nsp15/dsRNA | 6:2 nsp15/dsRNA |
|---|---|---|
| EMDB ID | 63618 | 63619 |
| PDB ID | 9M48 | 9M49 |
| **Data collection and processing** | | |
| Magnification | 105,000 | |
| Voltage (kV) | 300 | |
| Electron exposure (e–/Å$^2$) | 50 | |
| Defocus range (μm) | -1.5 to -1.8 | |
| Pixel size (Å) | 0.84 | |
| Symmetry imposed | C1 | C1 |
| Initial particle images (no.) | 319,161 | 319,161 |
| Final particle images (no.) | 244,604 | 117,130 |
| Map resolution (Å) | 2.6 | 2.78 |
| FSC threshold | 0.143 | 0.143 |
| Map resolution range (Å) | 2.9 to 6.1 | 3.2 to 6.7 |
| **Refinement** | | |
| Initial model used (PDB) | 7TJ2 | 9M48 |
| Model resolution (Å) | 2.8 | 3.2 |
| FSC threshold | 0.5 | 0.5 |
| Model resolution range (Å) | 50-2.8 | 50-3.2 |
| Map sharpening *B* factor (Å$^2$) | -67.2 | -67.2 |
| Model composition | | |
| Non-hydrogen atoms | 16977 | 18285 |
| Protein residues | 1989 | 1989 |
| Nucleotide residues | 62 | 124 |
| Ligands | 0 | 0 |
| *B* factors (Å$^2$) | | |
| Protein | 54.65 | 70.26 |
| Nucleotide | 21.48 | 27.67 |
| Ligand | 0 | 0 |
| R.m.s. deviations | | |
| Bond lengths (Å) | 0.002 | 0.002 |
| Bond angles (°) | 0.407 | 0.456 |
| Validation | | |
| MolProbity score | 1.38 | 1.36 |
| Clashscore | 4.5 | 5.41 |
| Poor rotamers (%) | 1.08 | 1.25 |
| Ramachandran plot | | |
| Favored (%) | 97.31 | 98.07 |
| Allowed (%) | 2.69 | 1.93 |
| Disallowed (%) | 0 | 0 |
